# Associations between sex, systemic iron and inflammatory status and subcortical brain iron

**DOI:** 10.1101/2023.08.22.554249

**Authors:** Holly Spence, Stephanie Mengoa-Fleming, Alan A. Sneddon, Christopher J. McNeil, Gordon D. Waiter

## Abstract

**Key Points:**

1. Differences in subcortical brain iron levels were observed between males and females
2. Females exhibited more associations between brain iron and systemic markers for iron than males.
3. Males exhibited more associations between brain iron and systemic markers for inflammation than females.

**Background:** Brain iron is increased in several neurodegenerative diseases, with studies demonstrating relationships between increased subcortical iron and disease progression. However, the causes of increased brain iron remain unclear. This study investigates relationships between subcortical iron and systemic iron and inflammatory status to improve our understanding of brain iron accumulation.

**Methods:** Brain MRI scans and blood plasma samples were collected from cognitively healthy females (n=176, mean age = 61.4 ± 4.5 y, age range = 28 - 72 y) andmales (n=152, mean age = 62.0 ± 5.1 y, age range = 32 - 74 y). Quantitative susceptibility mapping was used to quantify regional brain iron. To assess systemic iron, haematocrit, plasma ferritin and plasma soluble transferrin receptor (sTfR) were measured and total body iron index (TBI) was calculated. To assess systemic inflammation, C-reactive protein (CRP), Neutrophil/Lymphocyte ratio (NLR), plasma macrophage colony stimulating factor 1 (MCSF), plasma interleukin 6 (IL6) and plasma interleukin 1β (IL1β) were measured.

**Results:** Females exhibited associations between TBI and iron levels in the left and right caudate and right pallidum. Females also exhibited associations between haematocrit and iron levels in the right pallidum and left putamen. However, in males, the only association between brain iron and iron status was a positive correlation between iron in the right caudate and ferritin. In males, positive associations were observed between CRP levels and iron in the right thalamus and between IL6 levels and iron in the right amygdala and right pallidum. Males also exhibited a negative association between IL6 levels and iron in the left caudate. Positive associations between iron in the left thalamus and NLR were observed in both sexes.

**Conclusions:** This study demonstrates differential relationships between systemic iron and inflammation markers and brain iron in females and males. These results suggest differing iron regulation mechanisms in older age between sexes which could lead towards an understanding of the differences in neurodegenerative disease prevalence in males and females.

## Introduction

While brain iron is essential for many biological functions, increased deep grey matter iron has been observed in neurodegenerative diseases including Alzheimer’s disease (AD) and Parkinson’s disease (PD). Iron quantification via magnetic resonance imaging (MRI) may therefore be a promising tool for diagnosis of AD and PD (Kuchcinski, et al., 2023; Sato, et al., 2022; Lin, et al., 2023; Guan, et al., 2022; Liu, et al., 2021; Wang, et al., 2016). Several studies have also linked increases in brain iron observed in neurodegenerative disease with disease progression and cognitive decline (Yang, et al., 2022; Cogswell, et al., 2021; Spence, McNeil, & Waiter, 2020; Thomas, et al., 2020). Links between iron and neurodegeneration are thought to occur through the induction of oxidative stress and production of reactive oxygen species in Fenton’s reaction catalysed by iron and the subsequent induction of ferroptosis via interaction with lipid metabolism pathways (Wang, et al., 2022b; Floyd & Carney, 1992; Reichert, et al., 2020; Bertrand, 2017). In addition, imaging studies show that iron is found localised to protein aggregate plaques including Amyloid beta and Tau (Cogswell, et al., 2021; Everett, et al., 2020; Telling, et al., 2017; Fangola, Lee, Nixon, Duff, & Helpern, 2005). Iron has also been shown to interact with protein aggregates to enhance their aggregation and toxicity, while aggregated amyloid-β has also been shown to have a higher affinity for iron, contributing to links between iron and disease progression (Boopathi & Kolandaivel, 2016; Rottkamp, et al., 2001; Liu, et al., 2011; Guo, et al., 2013; Everett, et al., 2014). However, the exact mechanisms of increased brain iron accumulation are not yet clear.

Sternberg et al. observed increases in serum non-haem iron in AD, which correlate with cognitive measures of dementia, suggesting that brain iron accumulation associated with cognitive decline could occur due to a general increase in iron status (Sternberg, et al., 2017). However, relationships between brain iron and circulating markers of iron status are not well characterised, especially in the absence of disease. While some studies show positive correlations between brain iron and blood iron status, others demonstrate no association.(Gao, et al., 2017; Blasco, et al., 2017; Qiu, et al., 2014; Jakubowski, et al., 2021). Furthermore, dietary iron intake has been shown not to correlate with brain iron measured in an MRI study, suggesting that brain iron accumulation could occur independently of systemic iron levels (Valdes Hernandez M., et al., 2015).

Another potential mechanism regulating brain iron accumulation is through inflammatory pathways. Neuroinflammation has been associated with neurodegeneration via oxidative stress and can interact with iron homeostasis pathways via cytokines such as IL6 and IL1β (Guzman-Martinez, Maccioni, Andrade, & Patricio, 2019; Ganz, 2011; Smirnov, Bailey, Flowers, Garrigues, & Wesselius, 1999; Pinero, Hu, Cook, Scaduto Jr, & Connor, 2000). Increases in IL1β have been observed in AD, while increases in IL6 were observed in both AD and PD (Italiani, et al., 2018; Chaudhary, et al., 2021; Kwiatek-Majkusiak, et al., 2020). IL6 can directly activate iron regulatory hormone hepcidin, resulting in an increase in cellular iron levels through degradation and internalisation of the iron exporter ferroportin, blocking iron export (Urrutia, et al., 2013; Ganz, 2011). IL1β is known to interact with iron regulatory protein (IRP1) post-transcriptionally to increase its iron response element (IRE) binding capacity, resulting in increased ferritin translation, while treatment with IL1β was also shown to result in increased free iron uptake (Smirnov, Bailey, Flowers, Garrigues, & Wesselius, 1999; Pinero, Hu, Cook, Scaduto Jr, & Connor, 2000). Neuroinflammation has therefore been highlighted as a potential contributor towards increases in brain iron levels observed in neurodegenerative diseases.

This study aimed to examine the relationships between iron in the subcortical grey matter and systemic markers of iron and inflammatory status in order to better understand the mechanisms underlying iron accumulation in cognitively healthy individuals. To do this regional brain iron was quantified using quantitative susceptibility mapping and compared to blood markers of iron status (haematocrit, plasma ferritin and plasma soluble transferrin receptor (sTfR) and total body iron index (TBI) and markers of inflammatory status (C-reactive protein (CRP), Neutrophil: Lymphocyte ratio (NLR), plasma monoclonal colony stimulating factor 1 (MCSF), plasma interleukin 6 (IL6) and plasma interleukin 1β (IL1β). Due to the previously established differences in systemic iron regulation between sexes, primarily due to menstruation, pregnancy and menopause, in addition to the known sex differences in inflammatory cytokine levels due to hormonal regulation, males and females were investigated separately in this study (Whitfield, Treloar, Zhu, Powell, & Martin, 2003; Badenhorst, Forsyth, & Govus, 2022; Persson, et al., 2015).

## Methods

### Participants

In total, 176 females (mean age = 61.4 ± 4.5 years, age range = 28 - 72 years) and 152 males (mean age = 62.0 ± 5.1 years, age range = 32 - 74 years) were included in this study. Participants were recruited as part of the Stratifying Resilience and Depression Longitudinally (STRADL) study from the members of the Aberdeen Children of the 1950’s birth cohort (ACONF) and their first generation relatives. /this group took part in the Generation Scotland: Scottish Family Health Study (GS:SFHS) and had given consent to be recontacted, still lived in Scotland and had a community health index (CHI) number (Habota, et al., 2019). Ethical approval was previously obtained from the Scotland A Research Ethics Committee (REC reference number 14/55/0039) and the local Research and Development offices. Participants were excluded from the study if they did not have MRI data that was compatible with quantitative susceptibility mapping (QSM) processing, did not have sufficient volume of blood sample available for testing or had a neurodegenerative disease. In this study, the term “cognitively healthy” is used to signify lack of self-reported neurodegenerative disease. The age distribution of included participants is shown in Figure 1.

**Figure 1.**
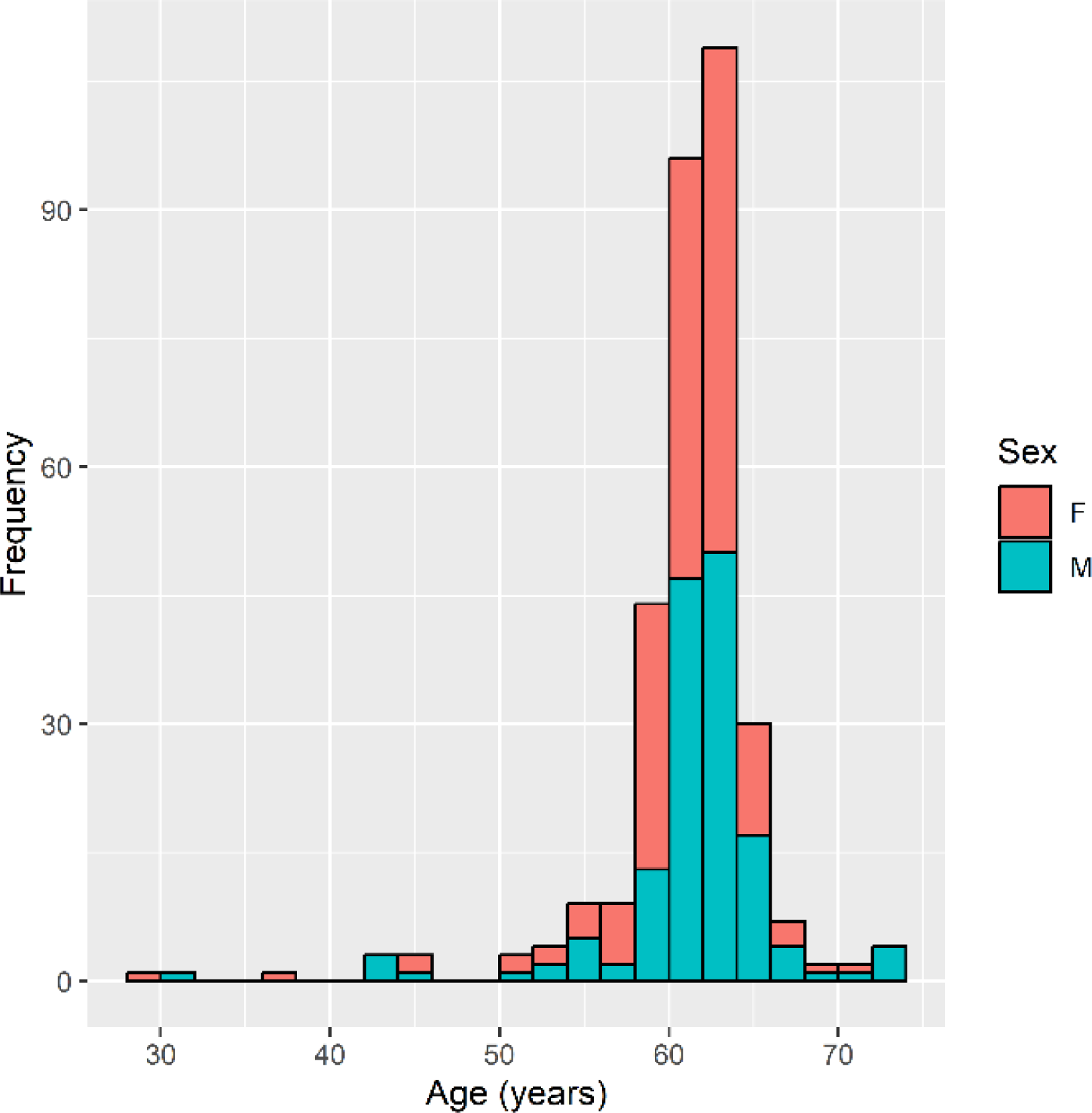
Histogram showing the age distribution of the included participants split by sex. F =Females (n=176), M = Males (n=152).

### Imaging

MRI protocol was previously described in full by Habota et al., 2019. For this study, T1 weighted, T2 weighted and multi-echo gradient-echo images (MEGRE) were used. All images were acquired on a 3T Philips Achieva TX-series MRI system (Philips Healthcare, Best, Netherlands) with a 32-channel phased-array head coil. 3D T1 weighted fast gradient echo images were acquired with the following parameters: 160 sagittal slices, TR = 8.2 ms, TE = 3.8 ms, TI = 1031 ms, FA = 8°, FOV = 240 mm, matrix size = 240 × 240, voxel size = 1.0 × 1.0 × 1.0 mm3, acquisition time = 5 min 38 s. T2 weighted images were acquired with the following parameters: 360 sagittal slices, TR = 2500 ms, TE = 314 ms, FA = 90°, FOV = 250 mm, matrix size = 252 × 250, voxel size = 0.5 × 0.5 × 0.5 mm3, acquisition time = 7 min 17 s. MEGRE images were acquired with the following parameters: 130 axial slices, TR = 31, TE = 7.2/13.4/19.6/25.8 ms, FA = 17°, FOV = 230 mm, matrix size = 384 x 316, Voxel size = 0.3 x 0.3 x 1 mm2, acquisition time = 4min 29s. Brain masks were generated using the brain extraction tool function of fsl software (Smith, 2022). Phase images from MEGRE data were processed through a QSM pipeline to calculate susceptibility maps used to estimate iron content as described previously (Spence, McNeil, & Waiter, 2022). Briefly, QSM was carried out using the STISuite V3.0 QSM GUI pipeline with Laplacian based phase unwrapping, a VSHARP background field correction and the iLSQR QSM method (Li, Wu, & Liu, 2013).

Susceptibility maps were co-registered with previously segmented T1 weighted images from which mean volume was calculated for the regions of interest (ROI), amygdala, caudate, hippocampus, pallidum, putamen, and thalamus (figure 2). Segmentation for these ROI was implemented using FreeSurfer software with the Desikan-Killiany atlas (Fischl, 2012). For hippocampal subfields cornu Ammonis (CA)1, CA3, CA4, subiculum and molecular layer of the hippocampus susceptibility maps were co-registered with T2 weighted images (with higher resolution than T1 weighted images) segmented using the previously described method from Iglesias et al., 2015 (Iglesias, et al., 2015). Susceptibility in the left and right hemispheres were calculated separately for all ROIs. As regional susceptibility data was skewed, regional median susceptibility rather than mean susceptibility was calculated for each ROI as a measure of regional iron content.

**Figure 2.**
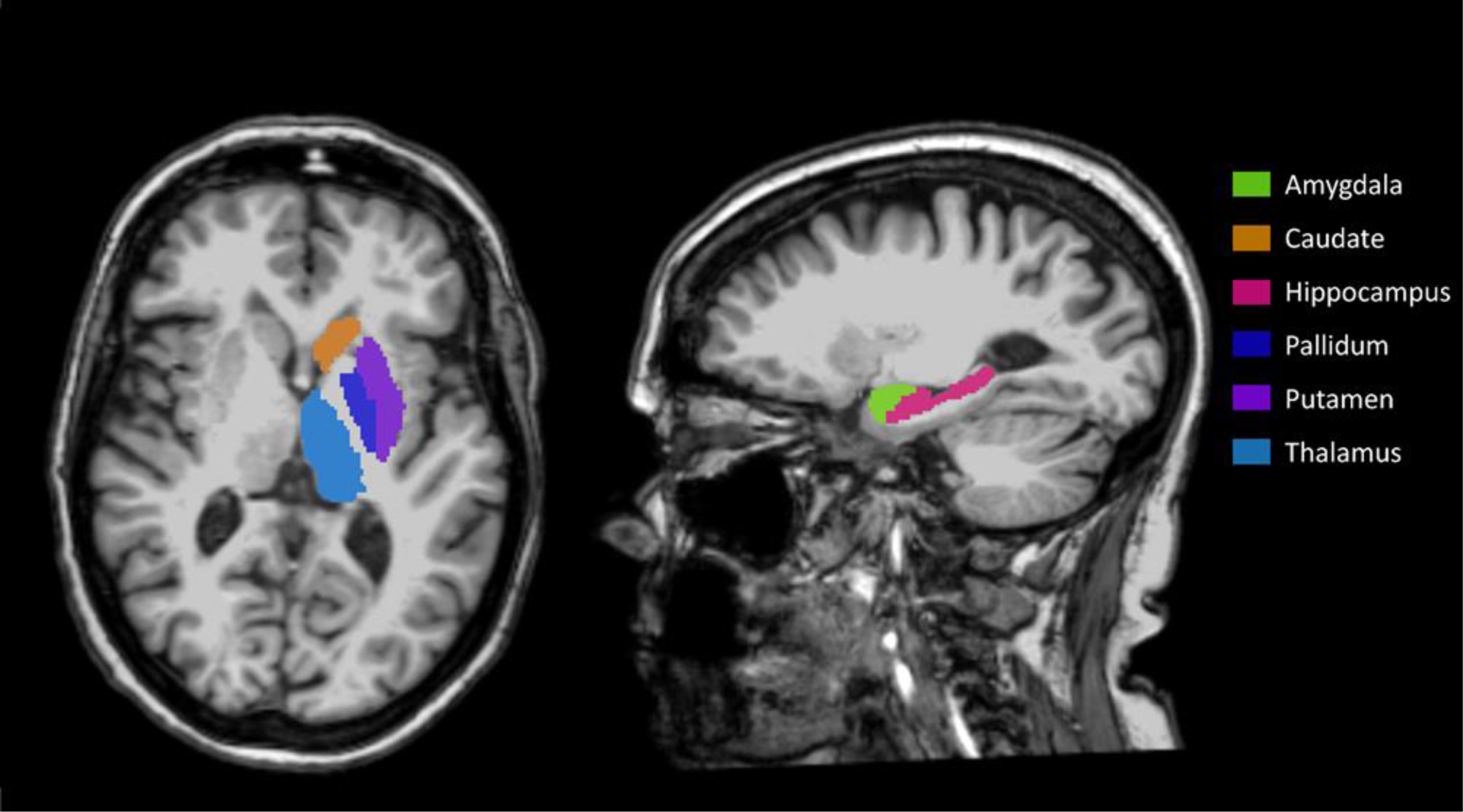
Segmented regions of interest implemented using FreeSurfer software with Desikan-Killiany atlas on T1 weighted image.

### Laboratory Analysis

Blood samples were taken from unfasted participants. Haematocrit, neutrophils, lymphocytes and CRP were analysed as part of a full blood count within NHS laboratories as described in the STRADL cohort profile (Habota, et al., 2019). CRP levels are described here as undetectable (<4 mg/L - below detection limit), normal (4-10 mg/L) or high (>10mg/L). NLR was calculated as neutrophil count/lymphocyte count.

Due to the differences in volumes of blood available per person, a different number of participants were included in each enzyme-linked immunosorbent assay (ELISA). 327 plasma samples were assessed for sTfR via an ELISA (RD194011100 - Oxford Biosystems Ltd UK). Absorbance was read at 450nm on a uQuant BioTek microplate spectrophotometer (Biotek Instruments, Inc). Plasma ferritin was assessed in 324 samples via an ELISA (RAB0197-1KT, Merck Life Science UK Ltd). TBI was calculated using the below equation as proposed by Cook et al., 2003 (Cook, Flowers, & Skikne, 2003):

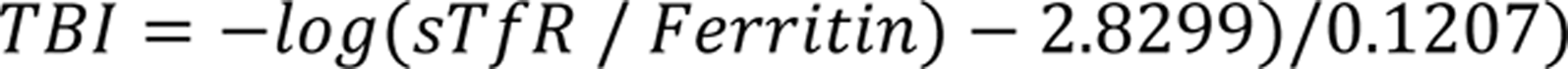

Plasma Macrophage colony-stimulating factor M-CSF was assessed via an ELISA in 325 participants (Authentikine KE00184, Proteintech Group, Inc). Plasma IL1β was assessed via ELISA in 320 participants (BMS224-2TEN – Thermo Fisher Scientific Inc.). Plasma IL6 was assessed via ELISA in 322 participants (BMS213-2TEN – Thermo Fisher Scientific Inc.). For inflammatory markers, absorbance was read at 450nm on an Infinite 200 PRO Tecan microplate reader.

### Statistical Analysis

All statistical analyses were carried out using R software (R Core Team, 2021). Pallidum iron was bimodally distributed and so was thresholded to generate a high and low iron group both with gaussian distribution. Susceptibility cut off points were set at 0.3 and 0.125 ppm for the left and right Pallidum respectively, based on the distribution of the data. No other regions were transformed for bimodal distribution. 14/322 participants had IL6 level below detection limit (DL) and so these values were imputed as DL/sqrt2, appropriate when the numbers of values below DL is <15%. IL6 was then treated as a continuous variable (Baccarelli, et al., 2005). 255/320 participants had IL1B levels below the DL and so IL1B data was split into two factors (above or below DL).

Before commencing formal analysis, it was determined that there was no significant difference in the age of females compared to males (p = 0.797). To assess differences between regional iron levels in males and females, variance between groups was first assessed. Where variance was equal, ANCOVA was used for parametric models where age and regional volume significantly contributed to the model as determined by Akaike information criterion (AIC). If age and regional volume did not contribute to the model, two sample t tests were used for parametric models. Welch’s t test was used for analysis of parametric models with unequal variance. To examine associations between blood markers and susceptibility, multivariate regression modelling was used where models were normally distributed and compared two continuous variables. Two sample t tests, chi squared, and ANCOVA were used for parametric models comparing groups where appropriate. For non-parametric distributions, data was log transformed into normality before further analysis. In all analysis, age, sex, volume, BMI and presence of infection were included as covariates where appropriate. AIC was used to determine whether confounding factors significantly contributed to the model. Variables were included if a decrease in AIC score was observed when the variable was added to the model, indicating less information was lost by the model. A Holm-Bonferroni multiple comparison correction was applied with a corrected α=0.05 denoting significance.

## Results

Table 1 summarises the demographics and blood marker data for the cohort.

**Table 1.** Cohort Demographics.

|  | Females | Males |
| --- | --- | --- |
| Age (yrs; mean ± SD) | 61.35 ± 4.52 | 61.96 ± 5.06 |
| BMI (mean ± SD) | 27.68 ± 5.81 | 27.22 ± 3.91 |
| Ferritin (ng/ml; mean ± SD) | 36.50 ± 40.13 | 51.19 ± 49.93 |
| sTfR (ug/ml; mean ± SD) | 0.97 ± 0.49 | 0.90 ± 0.41 |
| IL6 (pg/ml; mean ± SD) | 4.73 ± 5.92 | 4.89 ± 7.46 |
| MCSF (pg/ml; mean ± SD) | 588.64 ± 450.55 | 524.58 ± 363.27 |
| IL1β (Elevated:Undetectable) | 35:135 | 32:117 |
| CRP (High:Normal:Undetectable) | 0.478680556 | 0.135578704 |

### Differences in regional iron between sexes

Average median susceptibility for each ROI in males and females are shown in figure 4 and table 3. In whole grey matter (GM) ROIs, after correcting for multiple comparisons, males had significantly higher iron in the right hippocampus compared to females (p<0.001), while females had significantly higher iron in the right caudate compared to males (p=<0.001). Iron in males was shown to be higher in the left hippocampus (p=0.023), however this association did not remain significant after multiple comparison correction. Iron was shown to be higher in females in the left caudate (p=0.010), left putamen(p=0.044), left thalamus (p=0.046) and right pallidum (p=0.021), however none of these associations remained significant after correction for multiple comparison.

**Figure 3.**
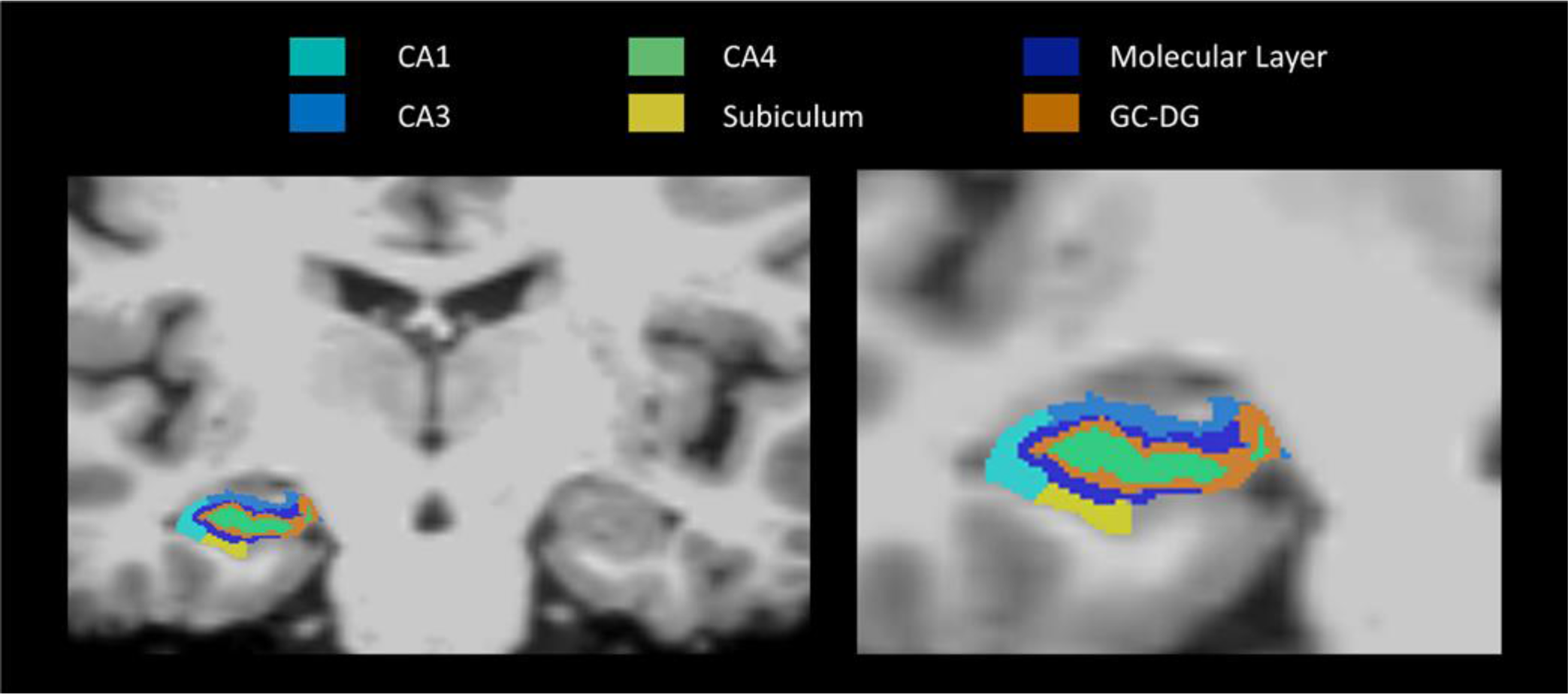
Hippocampal subfields segmented using FreeSurfer software with the Desikan-Killiany atlas shown here overlayed on a T1 weighted image. For this study, cornu Ammonis(CA)1, 3 and 4 were included as regions of interest, as well as the subiculum and molecular layer.

**Figure 4.**
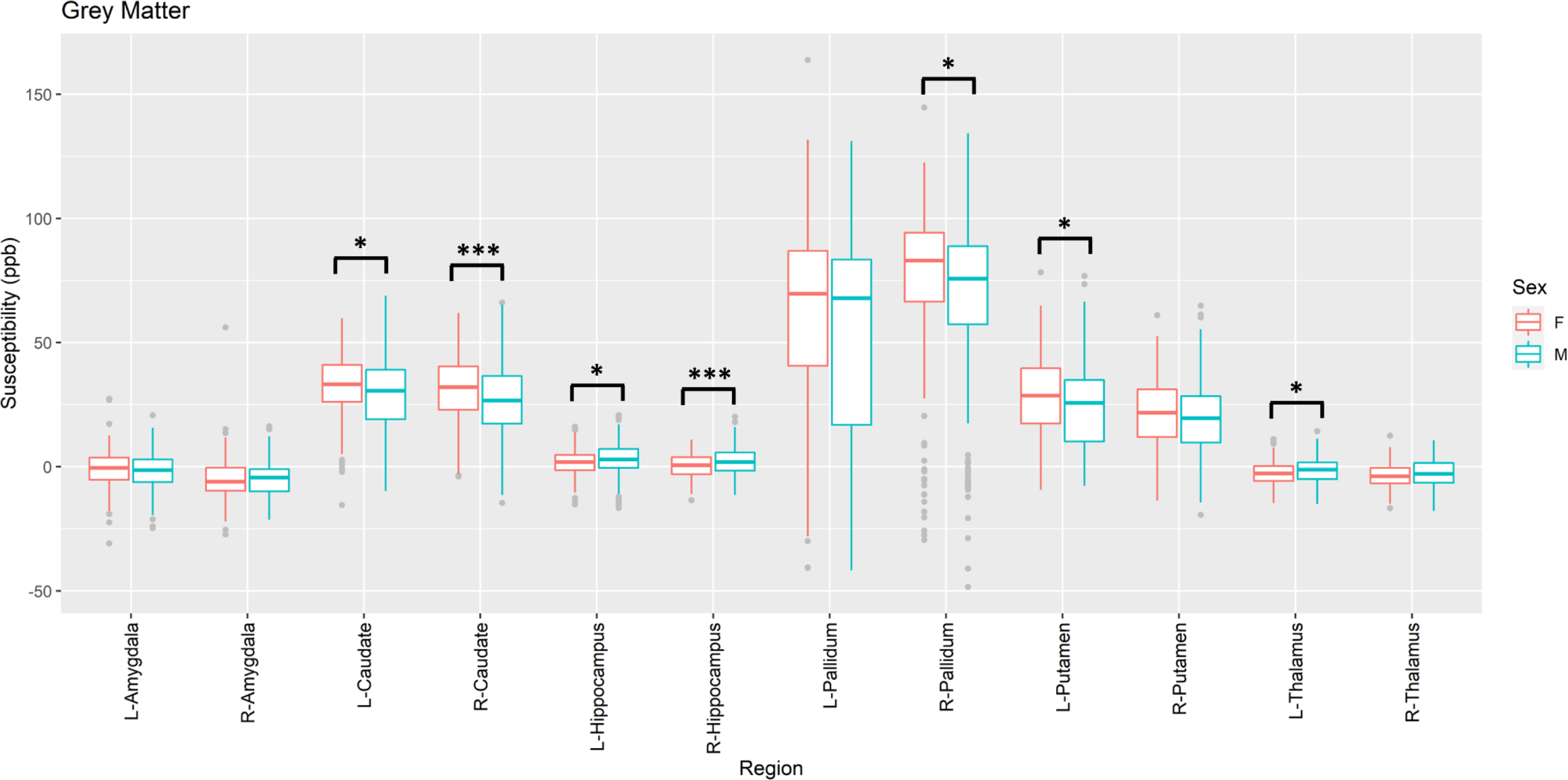
A) Sex differences in grey matter iron levels as determined by quantitative susceptibility mapping. * Indicates statistical significance where p<0.05, *** indicates statistical significance where p<0.001. F = Females (n=176), M = Males (n=152).

In the hippocampal subfields assessed, after multiple comparison correction, iron was significantly higher in males compared to females in the right CA1 (p<0.001), left and right CA3 (p<0.001; p<0.001), left and right CA4 (p<0.001; p<0.001), right subiculum (p=0.012) and right molecular layer (p<0.001). Iron in the left molecular layer was higher in males than females (p=0.029), but this result did not remain significant after correction for multiple comparison.

### Relationships between regional brain iron and blood iron status in females

Ferritin was positively associated with brain iron in the left and right pallidum in females (p=0.035, β=0.158, r2=0.051 and p=0.032, β=0.159, r2=0.069). Age and volume were included in the model for the left pallidum while only BMI was included in the right pallidum model as determined by AIC calculation. Total body iron index and haematocrit were also positively associated with right pallidum iron after controlling for BMI (p=0.015, β=0.180, r2=0.076 and p=0.013, β=0.184, r2=0.074 Total body iron index also positively associated with iron in the left and right caudate after controlling for BMI and volume or BMI only respectively (p=0.027, β=0.163, r2=0.079 and p=0.006, β=0.201, r2=0.086). Haematocrit was also positively associated with iron in the left putamen in females (p=0.021, β=0.174, r2=0.025). None of these associations between blood and brain iron status remained significant after multiple comparison correction. Blood iron status measures did not correlate with brain iron in any other regions in females. Relationships between regional brain iron and blood iron status markers in females are shown in Figure 5 *(supplementary figures 1 and 2)*.

**Figure 5.**
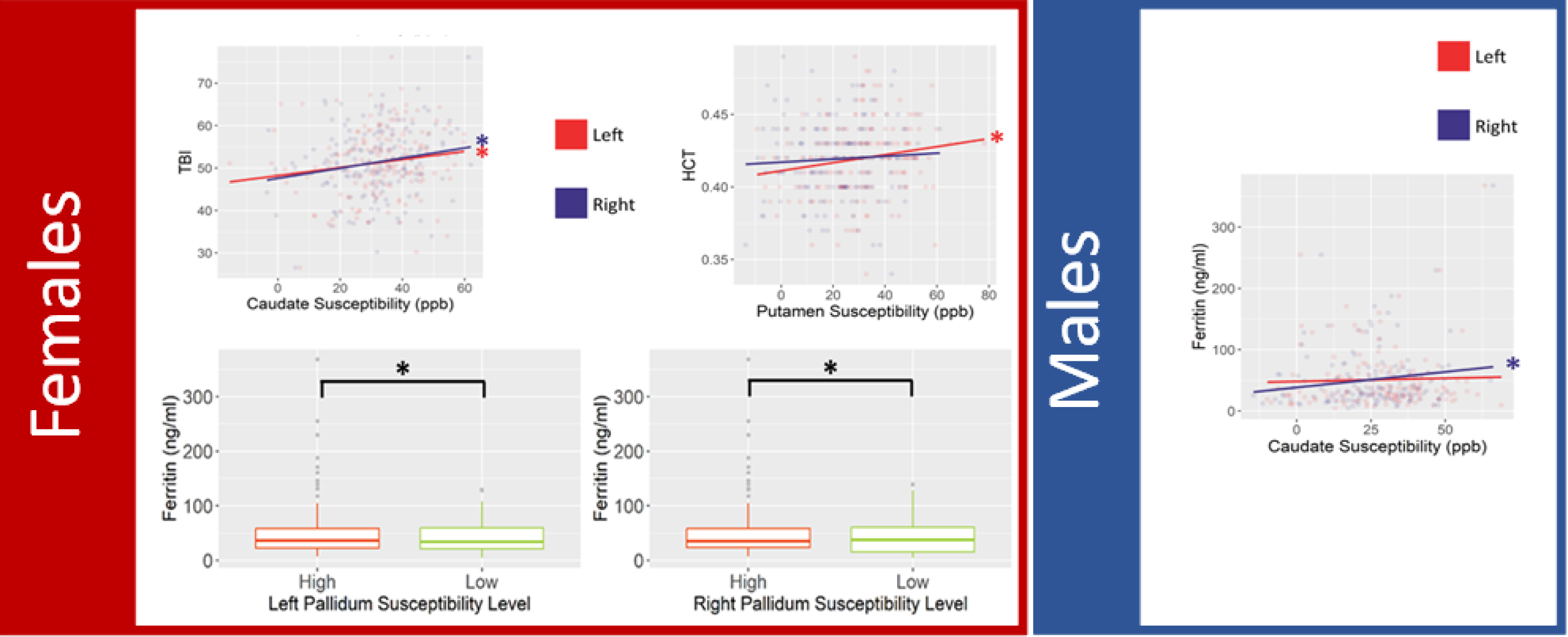
Statistically significant relationships between brain iron as measured by quantitative susceptibility mapping and markers for systemic iron, including Total body iron index (TBI), Haematocrit (HCT) and plasma ferritin were observed in females in the caudate, putamen and pallidum. Statistically significant relationships between brain iron as measured by quantitative susceptibility mapping and markers for systemic iron were only seen between caudate iron and ferritin in males. * Indicates statistical significance before multiple comparison correction where p<0.05.

### Relationships between regional brain iron and blood iron status in Males

Ferritin was positively associated with brain iron in the right caudate in males after controlling for BMI (p=0.041, β=0.168, r2=0.028). However, this association did not remain significant after multiple comparison correction and no other measures of blood iron were associated with brain iron in any other regions of interest in males. Relationships between regional brain iron and blood iron status markers in males are shown in Figure 5 *(supplementary figures 3 and 4)*.

### Relationships between regional brain iron and inflammatory status in females

In females, NLR was positively associated with iron in the left thalamus after controlling for age (p=0.034, β=0.160, r2=0.035). However, this association did not remain significant after multiple comparison correction and no other measures of inflammation were associated with brain iron in any other regions of interest in females. Relationships between regional brain iron and inflammatory status markers in females are shown in Figure 6 *(supplementary figures 5-8)*.

**Figure 6.**
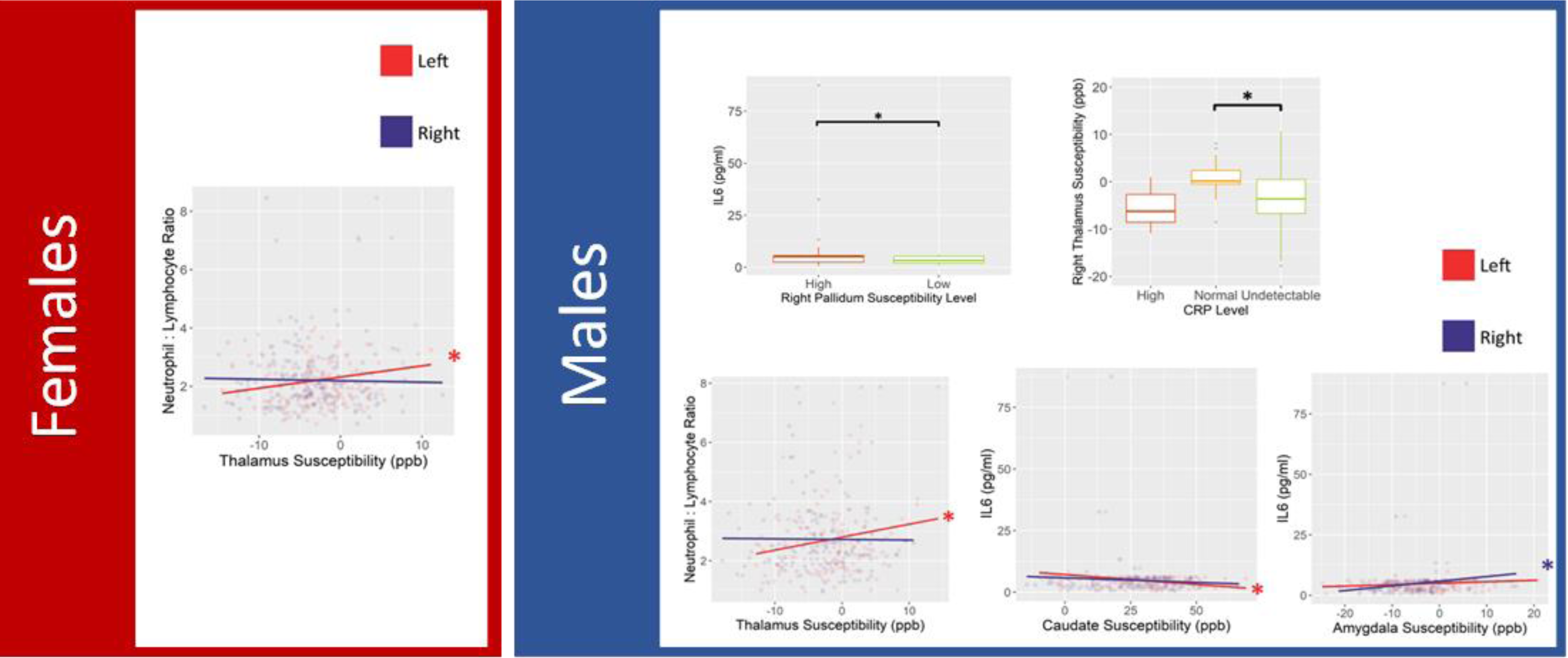
Statistically significant relationships between brain iron as measured by quantitative susceptibility mapping and markers for systemic inflammation were only observed between Neutrophil:Lymphocyte Ratio and Thalamus Susceptibility in females. Statistically significant relationships between brain iron as measured by quantitative susceptibility mapping and markers for systemic inflammation, including Neutrophil:Lymphocyte ratio, interleukin 6 (IL6) and C-reactive protein (CRP) were observed in the Pallidum, Thalamus, Caudate and Amygdala in males. * Indicates statistical significance before multiple comparison correction where p<0.05.

### Relationships between regional brain iron and inflammatory status in Males

In males, higher iron in the right thalamus was associated with normal CRP level compared to undetectable CRP (ANCOVA p = 0.039, post hoc p = 0.016). As in females, NLR was also positively associated with brain iron in the left thalamus (p=0.022, β=0.187, r2=0.029). Iron in the right amygdala was positively associated with IL6 (p=0.031, β=0.177, r2=0.025) and higher iron in the right pallidum was also associated with higher IL6 (p=0.038). However, after controlling for BMI and volume, IL6 was negatively associated with iron in the left caudate (p=0.025, β=-0.182, r2=0.047). Relationships between regional brain iron and inflammatory status markers in males are shown in Figure 6 *(supplementary figures 9-12)*.

Table 2 summarises the results of this study.

**Table 2.**
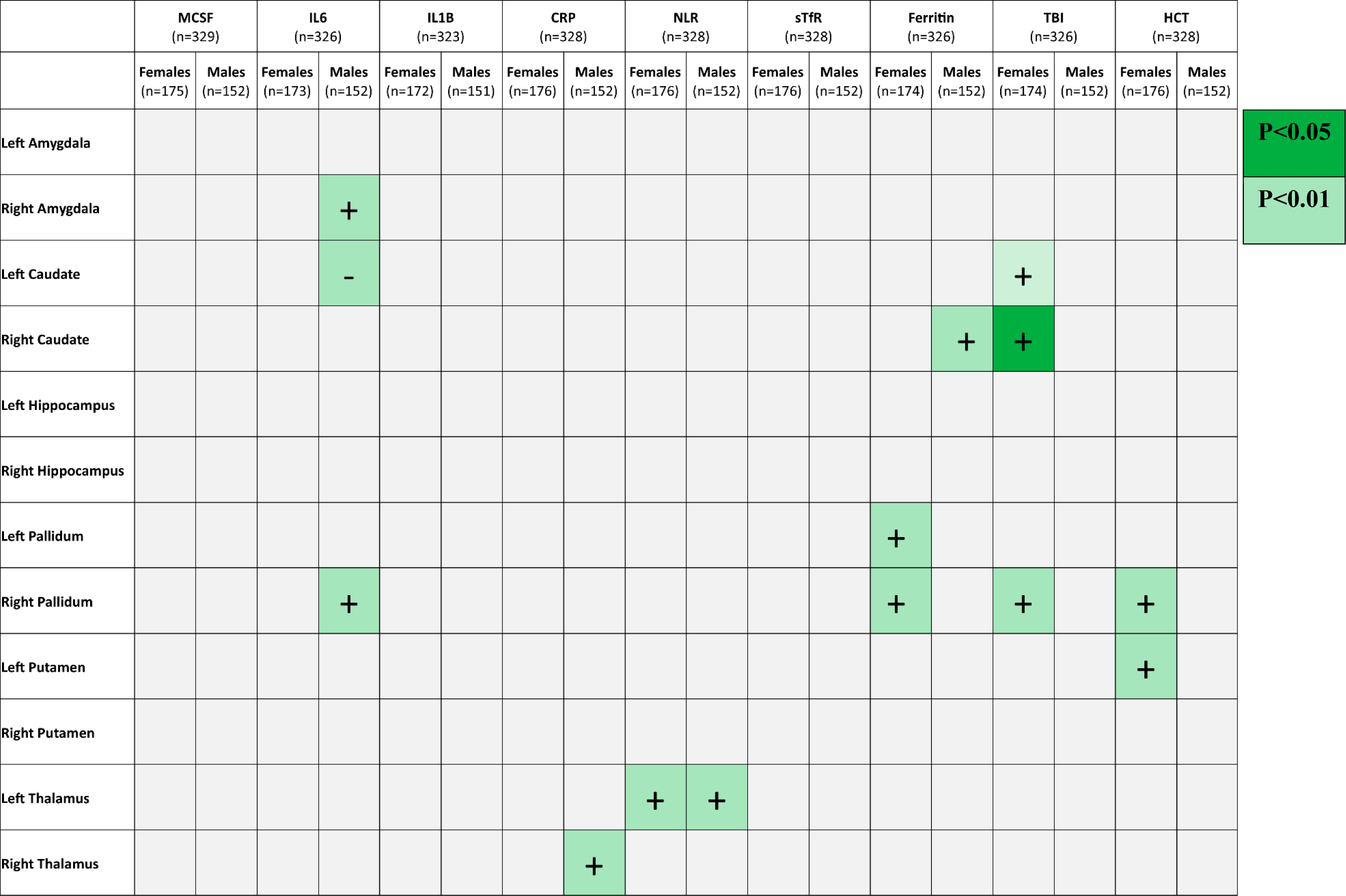
Summary table showing statistically significant correlations before multiple comparison correction between susceptibility in each ROI as measured by quantitative susceptibility mapping and blood markers for iron and inflammatory status. + indicates a positive correlation and – indicates a negative correlation.

## Discussion

Here, we demonstrated that females had lower iron in the right hippocampus compared to males, however, males had lower iron in the right caudate compared to females. When assessing relationships between brain iron and blood markers for iron and inflammation, several correlations were shown between brain iron and blood iron markers in females while, only one correlation was observed between brain iron and blood iron in males. However, several correlations were shown between blood inflammatory markers and brain iron in males, while only one correlation was shown in females. After multiple comparison correction, the only significant correlation between blood markers and brain iron was between NLR and iron in the left thalamus in both males and females.

This study demonstrated that in individuals without neurodegenerative disease, brain iron levels were lower in females compared to males in the right hippocampus. This observation adds to previous evidence for lower brain iron levels in females (Badenhorst, Forsyth, & Govus, 2022; Dugan, et al., 2021; Looker, Dallman, Carroll, Gunter, & Johnson, 1997). The reduced levels of iron in the hippocampus observed in females could be due to reduced iron availability due to the lasting effects of menstruation (Tishler, Raven, Lu, Altshuler, & Bartzokis, 2012). However, we also observed increased iron content in females in the right caudate region which would argue against this theory of menstruation related brain iron differences. It could be that due to the older mean age of the participants of this study, the majority of women here could be post-menopausal and thus no longer menstruating, which has been shown previously to impact brain iron levels (Tishler, Raven, Lu, Altshuler, & Bartzokis, 2012). These results suggests that different brain regions demonstrate different sex-mediated changes in iron regulation, in which the hippocampus appears to be more susceptible to sex-related changes in iron metabolism.

Before multiple comparison correction several associations between brain iron and systemic markers for iron and inflammatory status were significant, however, adjusting these findings using multiple comparison correction, resulted in several associations becoming non-significant. Therefore, conclusions about these associations must be made with caution, however, their biological plausibility and importance make their discussion justified.

In females, associations were found between higher blood iron measures and higher brain iron content across several regions including the pallidum, caudate and putamen. On the other hand, brain iron was not strongly associated with any blood iron markers in males, except for a weak association between ferritin and iron content in the right caudate. This suggests that brain iron in females is more impacted by blood iron levels, likely due to menstruation and higher anaemia incidence in women, whereas brain iron in males appears to be much less impacted by peripheral iron, if at all. The effects of the menopause are known to impact iron regulation lasting long after menopause ends (Tishler, Raven, Lu, Altshuler, & Bartzokis, 2012; Kim, Yetley, & Calvo, 1993). For example, changes in hepcidin that occur during exercise were shown to recover much slower in post-menopausal women which alters how effectively iron can be regulated by the body (Alfaro-Magallanes, et al., 2020). This means that blood iron status in females likely differs from males throughout life, initially due to menstruation and later due to the lasting effects of the menopause on sex hormone levels and iron regulation.

While only NLR was associated with left thalamus iron in females, males exhibited several positive correlations between inflammatory markers IL6, CRP and NLR and iron content in the amygdala, thalamus and pallidum. It is therefore likely that the mechanisms behind brain iron regulation and thus brain iron accumulation are different depending on sex. This may account for the difference in risk for neurodegenerative diseases associated with brain iron accumulation depending on sex. This is important for future research investigating the impact of brain iron on cognition and disease, as there is likely to be different causes behind these interactions depending on sex. Therefore, the efficacy of potential treatments for brain iron overload may differ between males and females. These results suggest that sex differences in the regulation of iron in the brain should be assessed separately rather than simply controlling for sex in multi-variate analysis.

A limitation of the current study is that systemic iron and inflammation levels were only measured in participants at a single time point. This lack of longitudinal measurement means that assessment of causation was not possible in this study. In the future, it is important for studies to assess the same participants over multiple time points, as it is possible that rate of increase in iron within an individual over time is a more appropriate measure. Longitudinal testing would also account for natural fluctuation in measures, for example, inflammatory status may be impacted by underlying infection. In addition, the markers measured here represent systemic circulating levels of inflammation and are not representative of local levels of inflammation in specific brain regions. It has been shown that some brain regions can exhibit targeted increases in pro-inflammatory cytokine expression, which can in turn have local effects on cell homeostasis (Smirnov, Bailey, Flowers, Garrigues, & Wesselius, 1999; Pinero, Hu, Cook, Scaduto Jr, & Connor, 2000; Urrutia, et al., 2013). Therefore, it could be that while significant relationships between brain iron and systemic inflammatory markers were not present, local levels of inflammation could still strongly impact brain iron.

A limitation of using QSM is that molecules other than iron also contribute to the signal measured including myelin. However, in this study, we only looked at myelin-poor grey matter regions where validation studies have shown that iron correlates directly with susceptibility (Hallgren & Sourander, 1958; Sun, et al., 2015; Langkammer, et al., 2012). It has been shown previously that post-menopausal women have lower brain iron deposition than men but higher blood iron than pre-menopausal women and so stage of menopause could account for some of the variation in this data (Kim, Nan, Kong, & Harlow, 2012; Persson, et al., 2015; Dugan, et al., 2021; Ofojekwu, et al., 2013). However, since most participants were over 55, it is likely that there was only a small portion of pre-menopausal participants and any variation in systemic iron due to menopause status or menstrual stage would be minimal. However, as participants did not disclose their menopause status as part of this study, it was not possible to look at pre-, peri- and post-menopausal groups separately.

Additionally, while it was determined that there was no difference in age between sexes in this study, there was a higher percentage of females than males. This is common in population-based recruitment where females tend to be more willing to volunteer than males. Volunteers also tend to be healthier and wealthier than the general population, which may also bias our findings.

## Conclusions

We demonstrate differences between males and females in associations between brain iron and systemic iron levels and also between brain iron and inflammatory status where we show that iron in the subcortical grey matter regions of males is more strongly associated with inflammatory status, whereas in females it is systemic iron status that is more strongly associated with brain iron than inflammatory status. Further study will be necessary to determine if the differences in significant results between males and females observed in this study are caused by an inherent biological difference in how brain iron is regulated between sexes. This is an important avenue of investigation as sex-related differences in brain iron could impact neurodegenerative disease risk and may impact research into efficacy therapeutic interventions targeting brain iron. These findings are also relevant for stratifying participants of trials for drugs such as iron chelators or inflammatory regulators, where males may respond better than females or vice versa. Further research into the mechanisms behind these sex differences may also show promise in understanding sex differences in risk for certain neurodegenerative diseases.

## Acknowledgments

We are grateful to the participants of the STRADL study which was supported and funded by the Wellcome Trust Strategic Award ‘Stratifying Resilience and Depression Longitudinally’ (STRADL) [104036/Z/14/Z]. This study was carried out as a part of Generation Scotland which received core support from the Chief Scientist Office of the Scottish Government Health Directorates [CZD/16/6] and the Scottish Funding Council [HR03006] and is currently supported by the Wellcome Trust [216767/Z/19/Z]. HS is supported by the Roland Sutton Academic Trust [0076/R/19]. We would also like to thank the STRADL project team.

## Data Availability Statement

The data collected in the STRADL study have been incorporated in the larger Generation Scotland dataset. Non-identifiable information from the Generation Scotland cohort is available to researchers in the United Kingdom and to international collaborators through application to the Generation Scotland Access Committee. Generation Scotland operates a managed data access process including an online application form, and proposals are reviewed by the Generation Scotland Access Committee.

## Author Contribution Statement

HS, AS, CM and GW contributed to the conception and design of the study. GW and HS performed the MRI data analysis. HS performed the plasma ELISA analysis. HS and SM-F performed subsequent statistical analysis and HS wrote the first draft of the manuscript. All authors contributed to manuscript revision, read and approved the submitted version.

## Abbreviations

AD: Alzheimer’s disease
AIC: Akaike information criterion
BET: Brain extraction tool
CRP: C-reactive protein
DL: Detection limit
M: Grey matter
HCT: Haematocrit
IL1β: Interleukin 1 beta
IL6: Interleukin 6
IRE: Iron response element
IRP: Iron regulatory protein
MCSF: Macrophage colony stimulating factor
MEGRE: Multi-echo gradient echo
MRI: Magnetic resonance imaging
NLR: Neutrophil:lymphocyte ratio
PD: Parkinson’s disease
QSM: Quantitative susceptibility mapping
SHARP: Sophisticated harmonic artifact reduction for phase data
sTfR: Soluble transferrin receptor
STRADL: Stratifying Resilience and Depression Longitudinally
TBI: Total body iron

