## SupplementaryTablesAndFigures for "Associations between sex, systemic iron and inflammatory status and subcortical brain iron"

Supplementary table 1. Average regional susceptibility as determined by quantitative susceptibility mapping in females and males.

| Region | Females (n=176) | | Males (n=152) | | p-value |
|---|---|---|---|---|---|
| | Mean Susceptibility (ppb) | Standard Deviation (ppb) | Mean Susceptibility (ppb) | Standard Deviation (ppb) | |
| Left Amygdala | -1.170 | 8.162 | -1.728 | 7.690 | |
| Right Amygdala | -5.047 | 8.280 | -5.028 | 6.689 | |
| Left Caudate | 32.525 | 13.070 | 28.664 | 15.883 | |
| Right Caudate | 31.297 | 12.946 | 26.099 | 15.164 | |
| Left Hippocampus | 1.707 | 5.192 | 3.088 | 6.633 | |
| Right Hippocampus | 0.492 | 4.916 | 2.347 | 5.723 | |
| Left Pallidum | 60.567 | 38.414 | 52.599 | 42.484 | |
| Right Pallidum | 75.731 | 31.159 | 67.808 | 34.621 | |
| Left Putamen | 28.360 | 16.059 | 24.362 | 17.529 | |
| Right Putamen | 21.646 | 14.693 | 19.208 | 14.836 | |
| Left Thalamus | -2.665 | 4.604 | -1.629 | 5.334 | |
| Right Thalamus | -3.397 | 5.076 | -2.756 | 5.835 | |
| Left CA1 | 2.502 | 8.102 | 3.899 | 9.279 | |
| Right CA1 | 1.551 | 6.612 | 4.290 | 7.836 | |
| Left CA3 | -5.773 | 9.138 | -2.122 | 10.633 | |
| Right CA3 | -1.278 | 9.661 | 3.241 | 10.798 | |
| Left CA4 | -5.286 | 9.658 | -1.805 | 10.407 | |
| Right CA4 | -5.910 | 9.941 | -0.982 | 10.587 | |
| Left Molecular Layer | 4.465 | 7.263 | 6.160 | 8.386 | |
| Right Molecular Layer | 1.446 | 8.104 | 6.195 | 7.685 | |
| Left Subiculum | 7.388 | 6.580 | 8.082 | 9.296 | |
| Right Subiculum | 2.476 | 6.458 | 3.787 | 9.744 | |

### Females

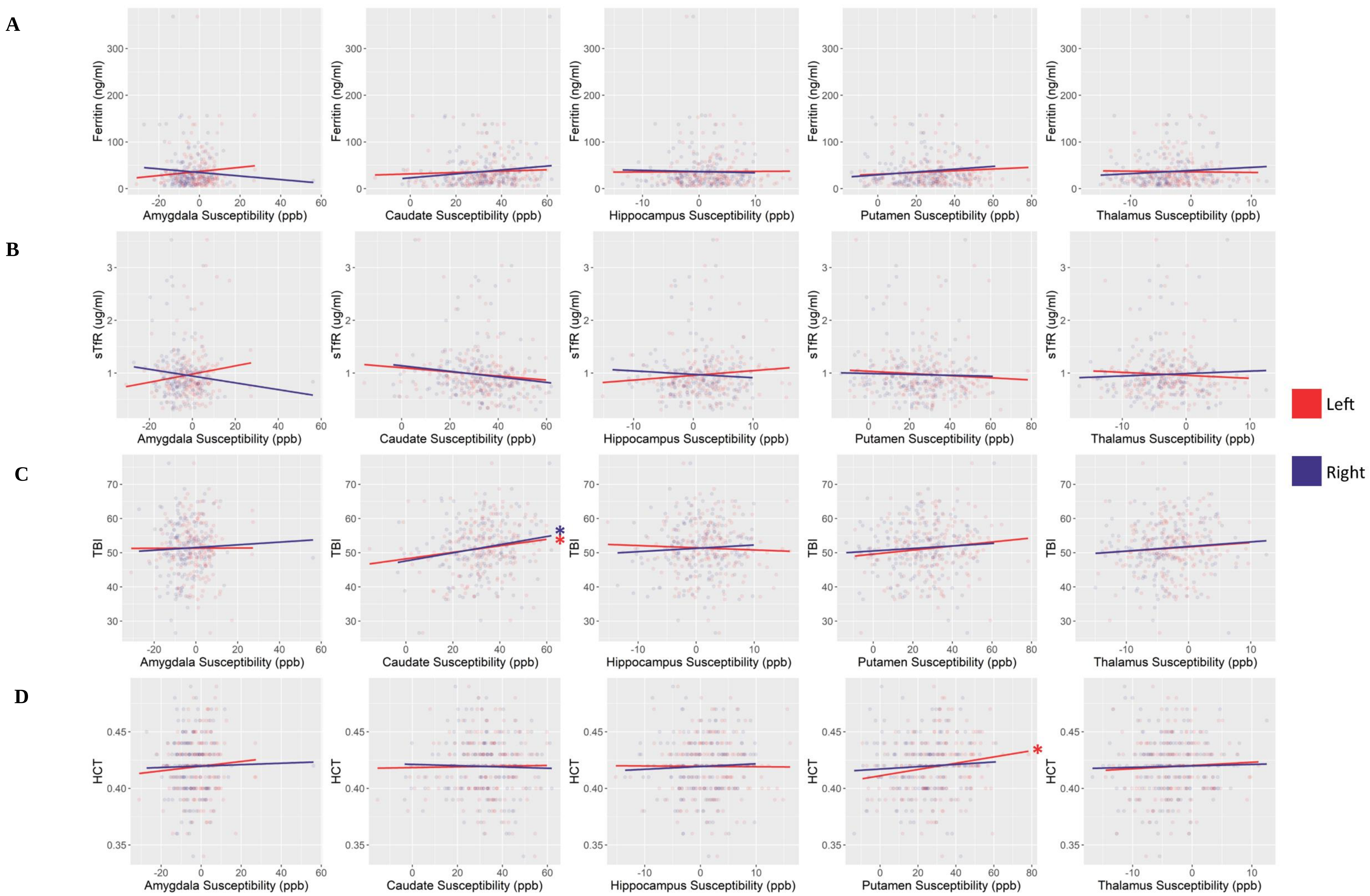

A
B
C
D
Ferritin (ng/ml)
sTfR (ug/ml)
TBI
HCT
Amygdala Susceptibility (ppb)
Caudate Susceptibility (ppb)
Hippocampus Susceptibility (ppb)
Putamen Susceptibility (ppb)
Thalamus Susceptibility (ppb)
*
Left
Right

Supplementary Figure 1. A) Relationships in females between plasma ferritin and brain iron in grey matter regions as measured by quantitative susceptibility mapping. (n=174). B) Relationships in females between plasma soluble transferrin receptor (sTfR) and brain iron in grey matter regions as measured by quantitative susceptibility mapping. (n=176). C) Relationships in females between total body iron (TBI) index and brain iron in grey matter regions as measured by quantitative susceptibility mapping. (n=174). D) Relationships in females between haematocrit (HCT) and brain iron in grey matter regions as measured by quantitative susceptibility mapping. (n=176). * Indicates statistical significance before multiple comparison correction where $p<0.05$.

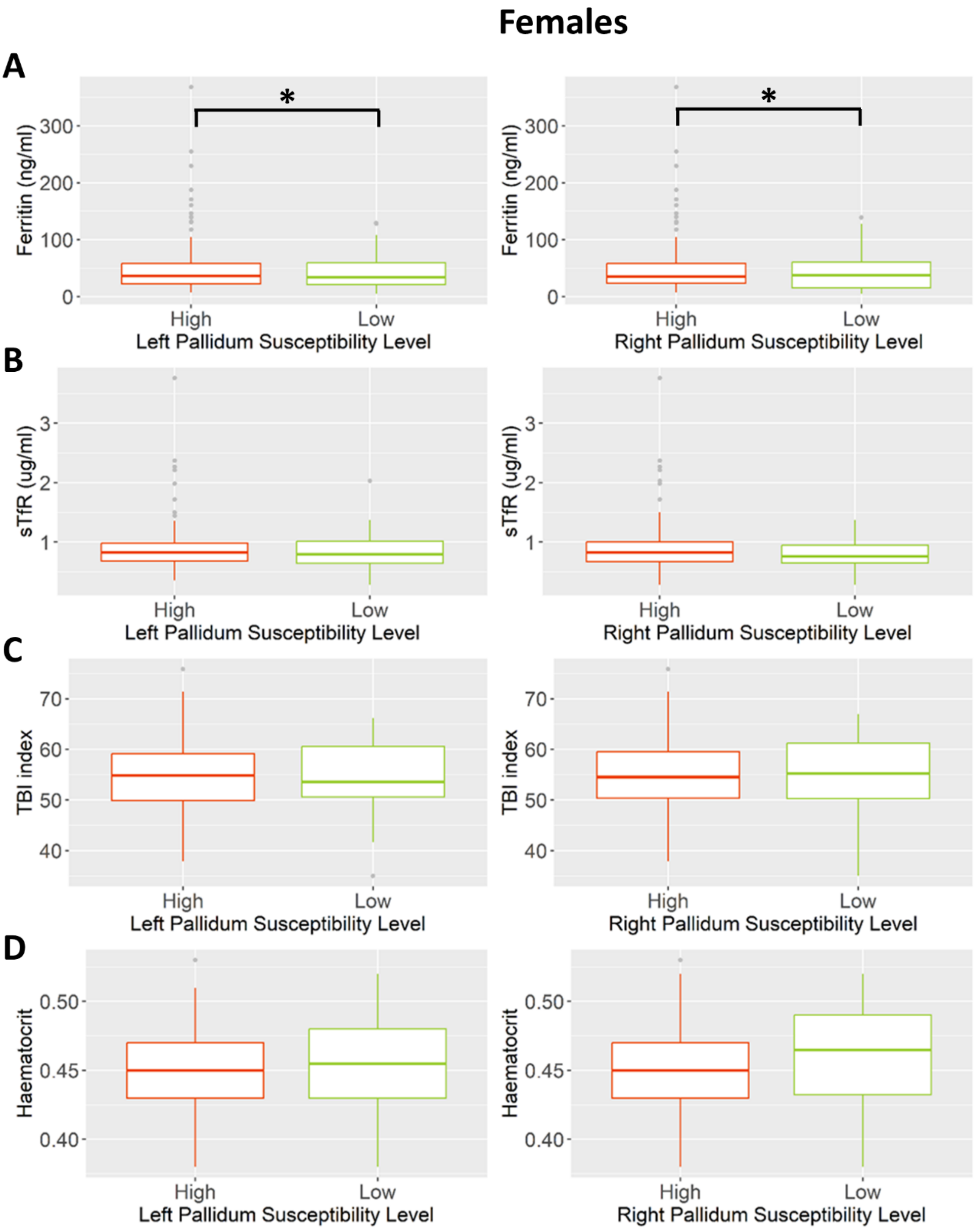


Supplementary Figure 2. A) Relationships in females between plasma ferritin and brain iron in the pallidum as measured by quantitative susceptibility mapping. (n=174). B) Relationships in females between log(plasma soluble transferrin receptor (sTfR)) and brain iron in the pallidum as measured by quantitative susceptibility mapping. (n=176). C) Relationships in females between total body iron (TBI) index and brain iron in the pallidum as measured by quantitative susceptibility mapping. (n=174). D) Relationships in females between haematocrit and brain iron in the pallidum as measured by quantitative susceptibility mapping. (n=176). High = left pallidum susceptibility > 0.3ppm OR right pallidum susceptibility >0.125ppm, Low = left pallidum susceptibility < 0.3ppm OR right pallidum susceptibility < 0.125ppm. * Indicates statistical significance before multiple comparison correction where $p<0.05$.

### Males

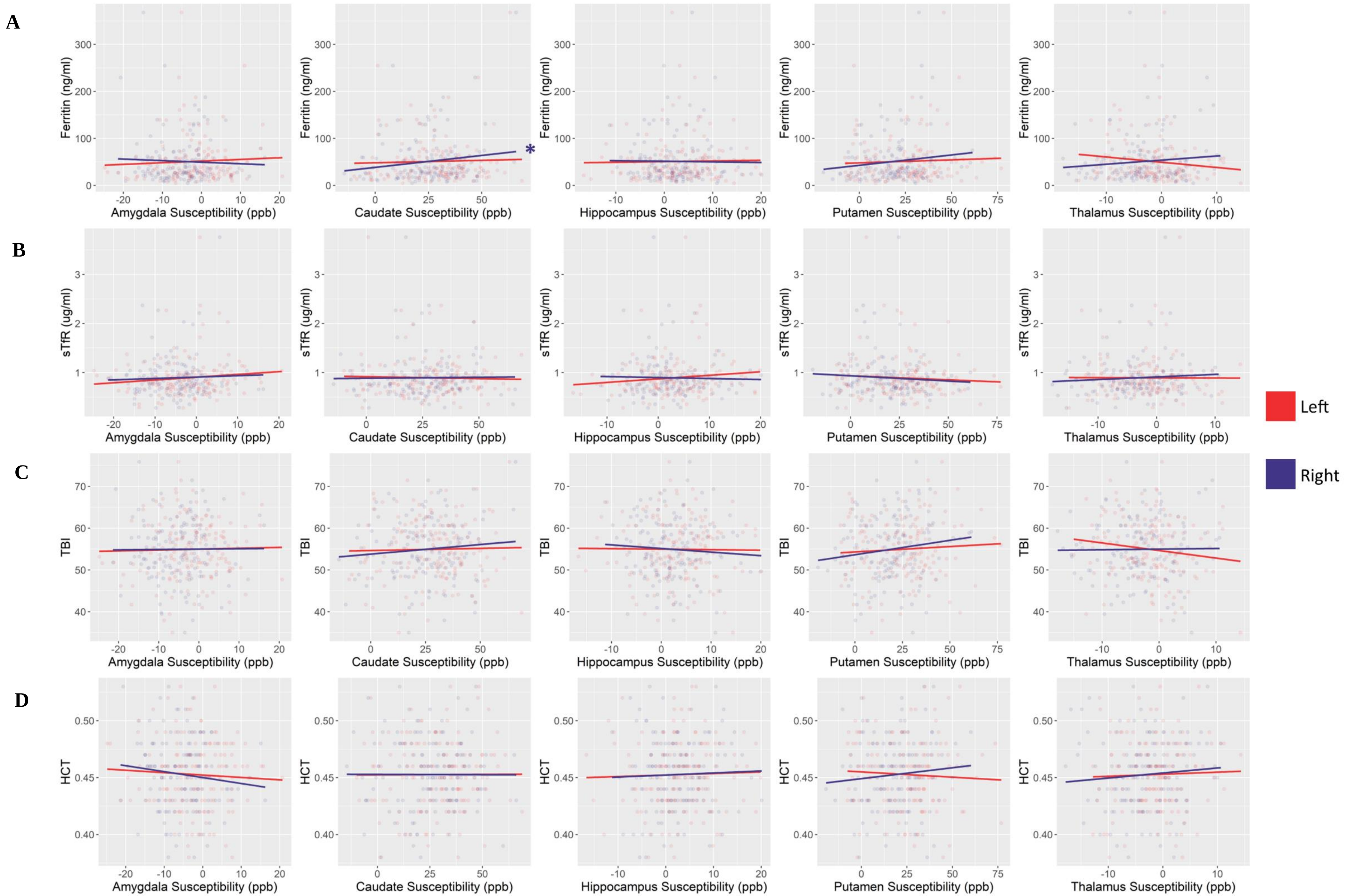

A
B
C
D
Ferritin (ng/ml)
sTfR (ug/ml)
TBI
HCT
Amygdala Susceptibility (ppb)
Caudate Susceptibility (ppb)
Hippocampus Susceptibility (ppb)
Putamen Susceptibility (ppb)
Thalamus Susceptibility (ppb)
*
Left
Right

Supplementary Figure 3. A) Relationships in males between plasma ferritin and brain iron in grey matter regions as measured by quantitative susceptibility mapping. (n=152). B) Relationships in males between plasma soluble transferrin receptor (sTfR) and brain iron in grey matter regions as measured by quantitative susceptibility mapping. (n=152). C) Relationships in males between total body iron (TBI) index and brain iron in grey matter regions as measured by quantitative susceptibility mapping. (n=152). D) Relationships in males between haematocrit and brain iron in grey matter regions as measured by quantitative susceptibility mapping. (n=152). * Indicates statistical significance before multiple comparison correction where $p<0.05$

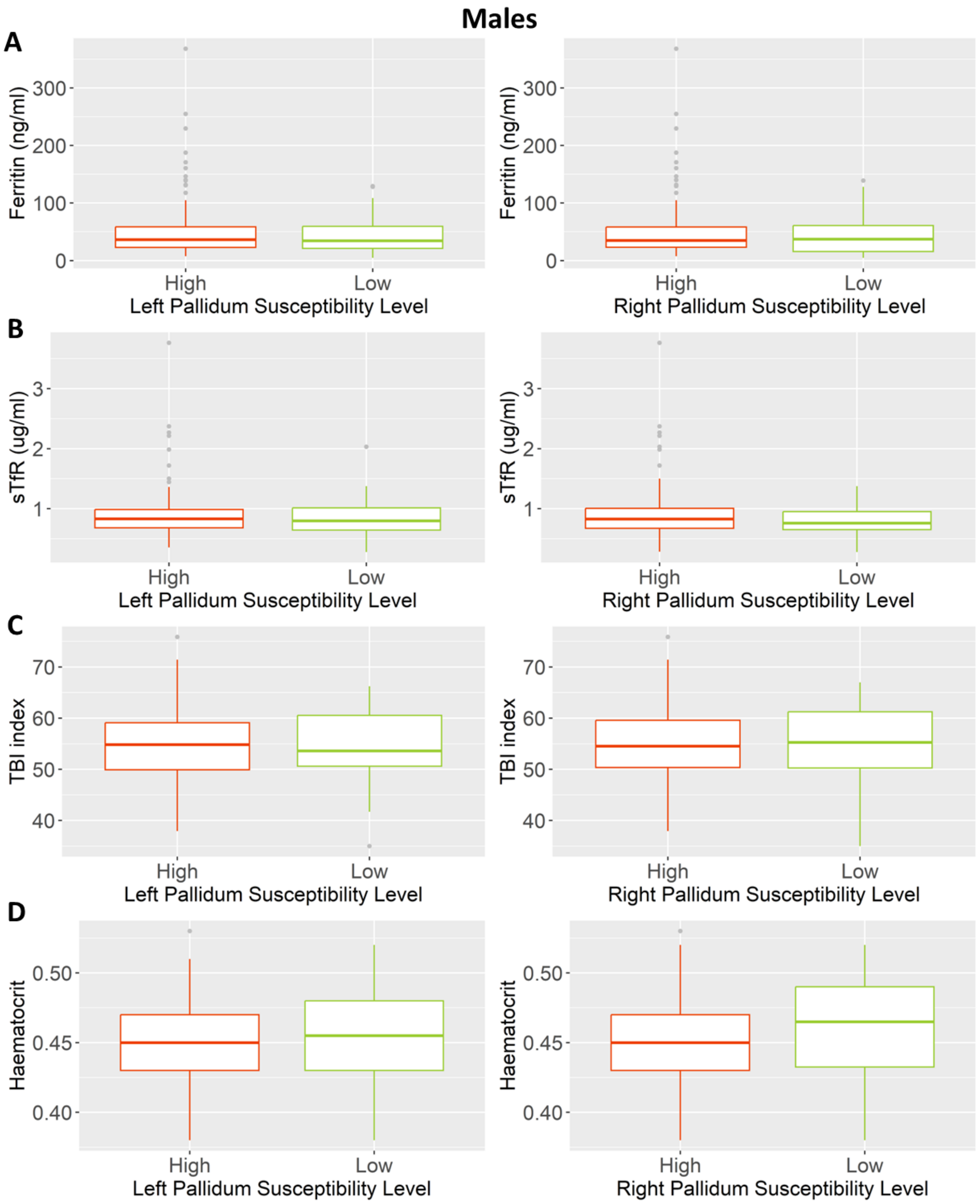


Supplementary Figure 4. A) Relationships in males between plasma ferritin and brain iron in the pallidum as measured by quantitative susceptibility mapping. (n=152). B) Relationships in males between plasma soluble transferrin receptor (sTfR) and brain iron in the pallidum as measured by quantitative susceptibility mapping. (n=152). C) Relationships in males between total body iron (TBI) index and brain iron in the pallidum as measured by quantitative susceptibility mapping. (n=152). D) Relationships in males between haematocrit and brain iron in the pallidum as measured by quantitative susceptibility mapping. (n=152). High = left pallidum susceptibility > 0.3ppm OR right pallidum susceptibility >0.125ppm, Low = left pallidum susceptibility < 0.3ppm OR right pallidum susceptibility < 0.125ppm.

### Females

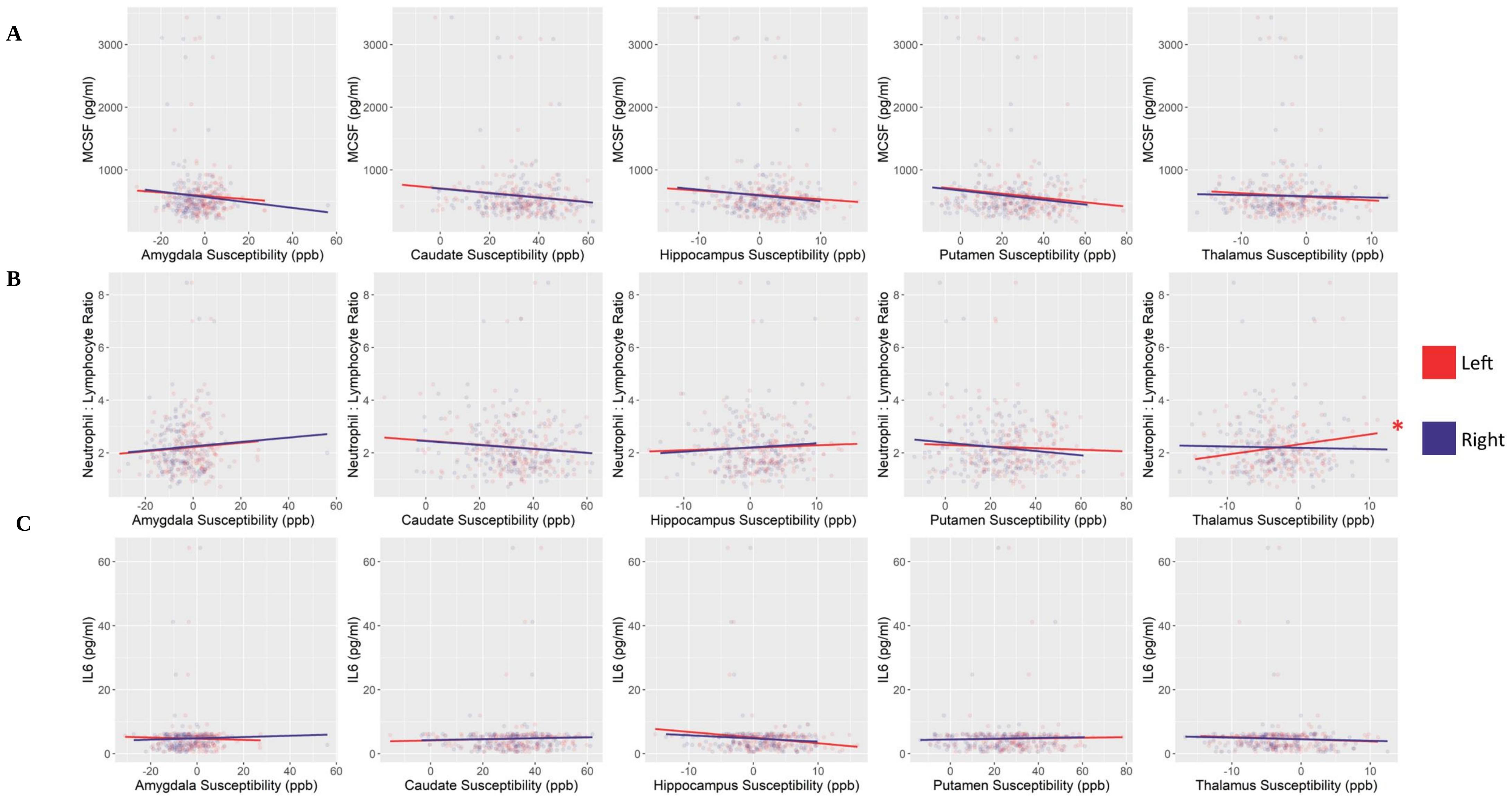

A
B
C
MCSF (pg/ml)
Neutrophil : Lymphocyte Ratio
IL6 (pg/ml)
Amygdala Susceptibility (ppb)
Caudate Susceptibility (ppb)
Hippocampus Susceptibility (ppb)
Putamen Susceptibility (ppb)
Thalamus Susceptibility (ppb)
Left
Right
*

Supplementary Figure 5. A) Relationships in females between plasma monoclonal colony stimulating factor 1 (MCSF) and brain iron in grey matter regions as measured by quantitative susceptibility mapping. (n=175). B) Relationships in females between neutrophil lymphocyte ratio and brain iron in grey matter regions as measured by quantitative susceptibility mapping. (n=176). C) Relationships in females between plasma interleukin – 6 (IL6) and brain iron in grey matter regions as measured by quantitative susceptibility mapping. (n=173). * Indicates statistical significance before multiple comparison correction where $p<0.05$.

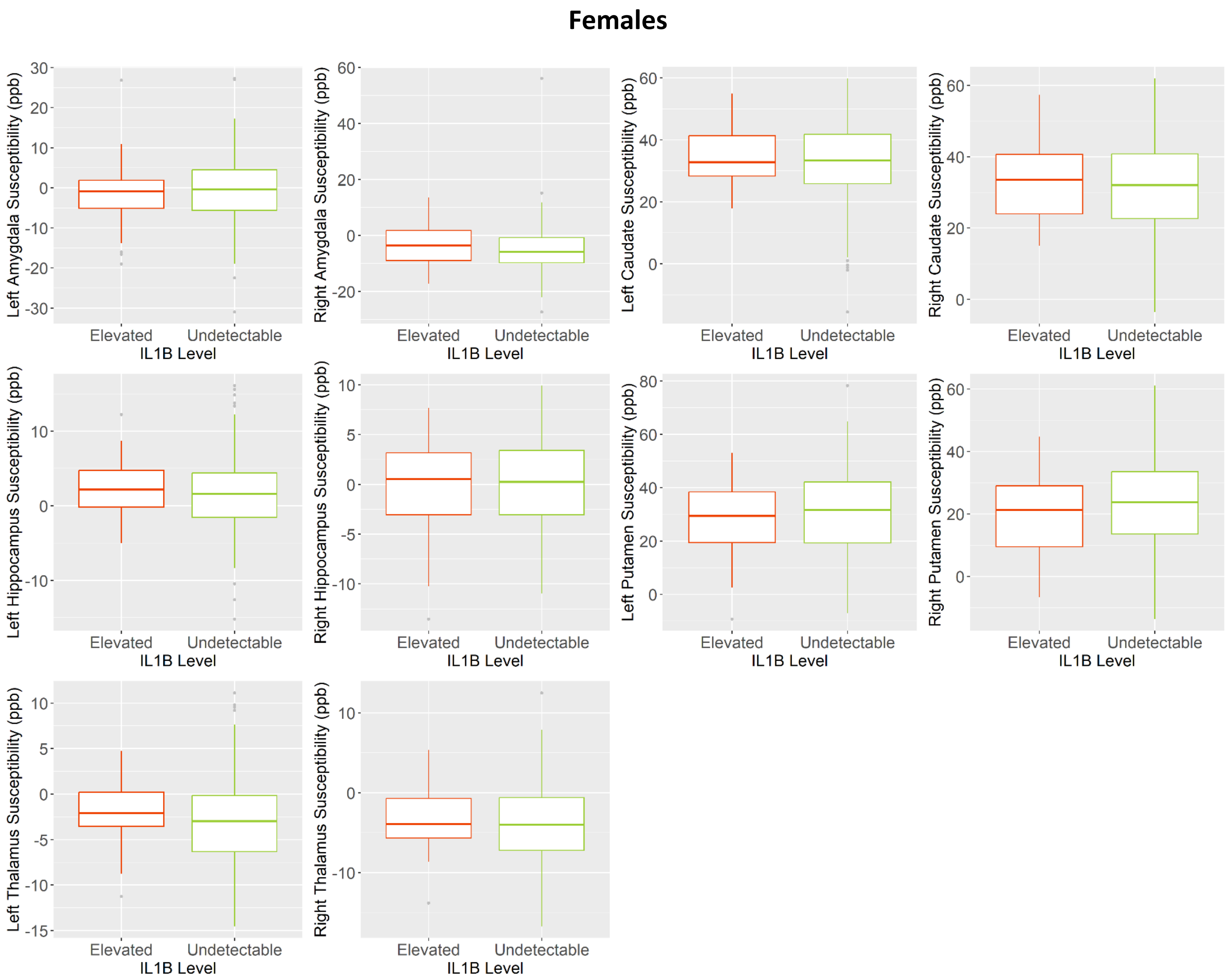
Females
Left Amygdala Susceptibility (ppb)
Right Amygdala Susceptibility (ppb)
Left Caudate Susceptibility (ppb)
Right Caudate Susceptibility (ppb)
Left Hippocampus Susceptibility (ppb)
Right Hippocampus Susceptibility (ppb)
Left Putamen Susceptibility (ppb)
Right Putamen Susceptibility (ppb)
Left Thalamus Susceptibility (ppb)
Right Thalamus Susceptibility (ppb)
Elevated
Undetectable
IL1B Level

Supplementary Figure 6. Box and whisker plots showing the relationships in females between plasma interleukin 1 beta (IL1B) and iron in grey matter regions as measured by quantitative susceptibility mapping (n=172). Elevated = above detection limit (>0.92pg/ml), Undetectable = below detection limit (<0.92pg/ml).

### Females

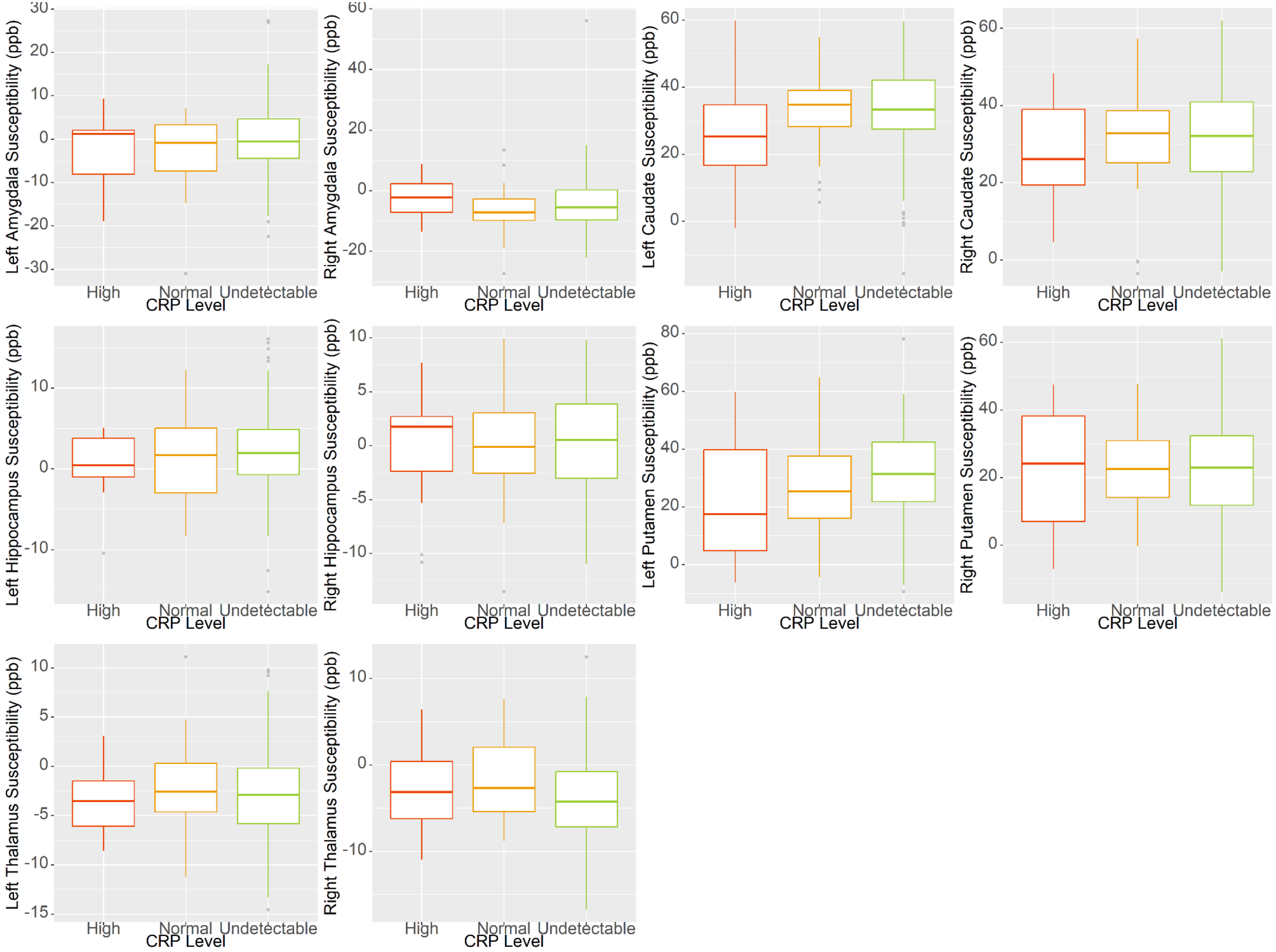

Left Amygdala Susceptibility (ppb)
Right Amygdala Susceptibility (ppb)
Left Caudate Susceptibility (ppb)
Right Caudate Susceptibility (ppb)
Left Hippocampus Susceptibility (ppb)
Right Hippocampus Susceptibility (ppb)
Left Putamen Susceptibility (ppb)
Right Putamen Susceptibility (ppb)
Left Thalamus Susceptibility (ppb)
Right Thalamus Susceptibility (ppb)
High
Normal
Undetectable
CRP Level

Supplementary Figure 7. Box and whisker plots showing relationships in females between c-reactive protein (CRP) and iron in grey matter regions as measured by quantitative susceptibility mapping (n=176). High = >10mg/L, normal = 4-10mg/L, <4mg/L. * Indicates statistical significance before multiple comparison correction

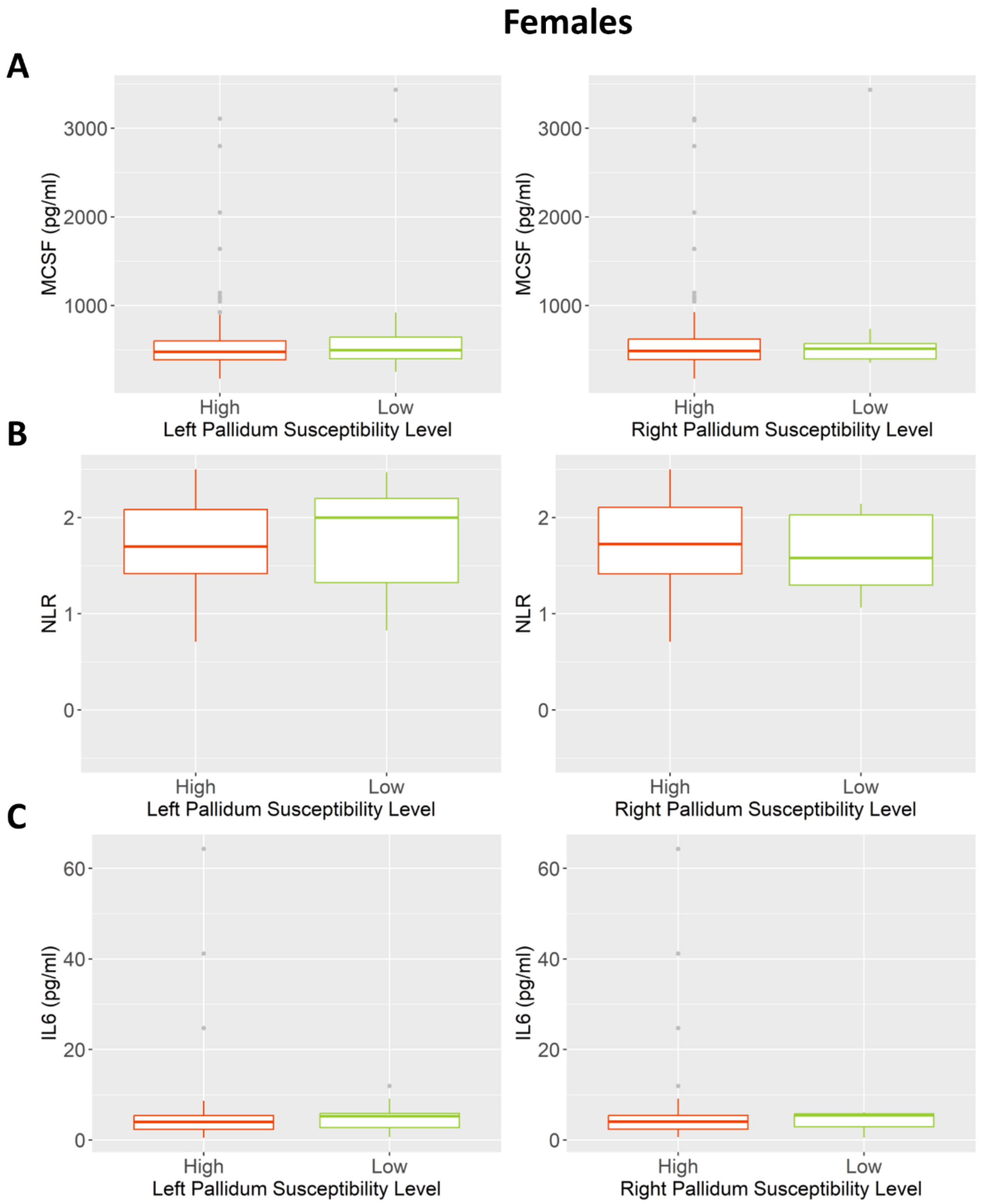


Supplementary Figure 8. A) Relationships in females between plasma macrophage colony stimulating factor 1 (MCSF) and brain iron in the pallidum as measured by quantitative susceptibility mapping (n=175). B) Relationships in females between neutrophil lymphocyte ratio and brain iron in the pallidum as measured by quantitative susceptibility mapping (n=176). C) Relationships in females between plasma interleukin – 6 (IL6) and brain iron in the pallidum as measured by quantitative susceptibility mapping (n=173). High = left pallidum susceptibility > 0.3ppm OR right pallidum susceptibility >0.125ppm, Low = left pallidum susceptibility < 0.3ppm OR right pallidum susceptibility < 0.125ppm.

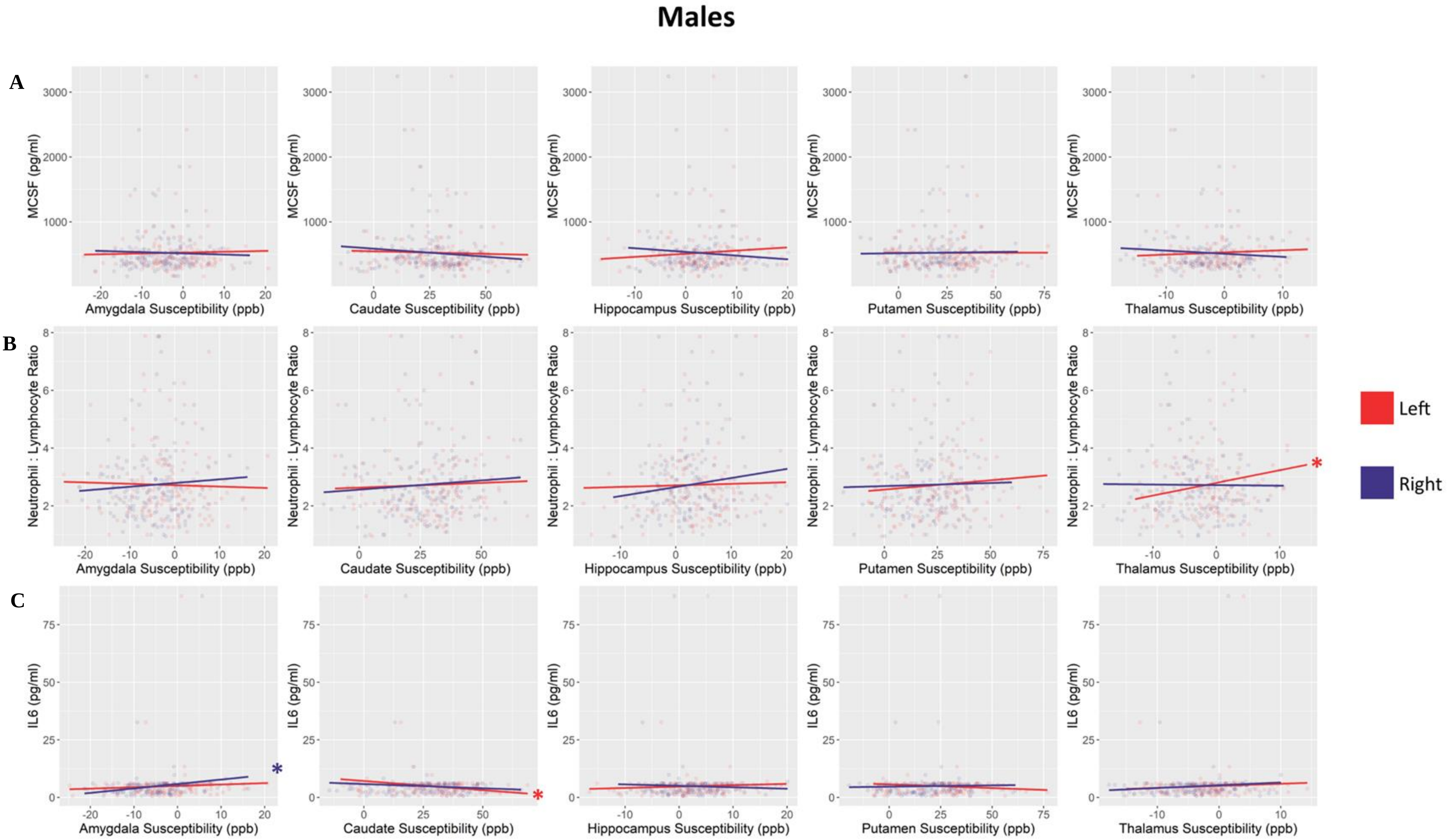
Males
A
B
C
MCSF (pg/ml)
Neutrophil : Lymphocyte Ratio
IL6 (pg/ml)
Amygdala Susceptibility (ppb)
Caudate Susceptibility (ppb)
Hippocampus Susceptibility (ppb)
Putamen Susceptibility (ppb)
Thalamus Susceptibility (ppb)
Left
Right

Supplementary Figure 9. A) Relationships in males between plasma macrophage colony stimulating factor 1 (MCSF) and brain iron in grey matter regions as measured by quantitative susceptibility mapping (n=154). B) Relationships in males between neutrophil lymphocyte ratio and brain iron in grey matter regions as measured by quantitative susceptibility mapping (n=155). C) Relationships in males between plasma interleukin – 6 (IL6) and brain iron in grey matter regions as measured by quantitative susceptibility mapping (n=153). * Indicates statistical significance before multiple comparison correction where $p<0.05$.

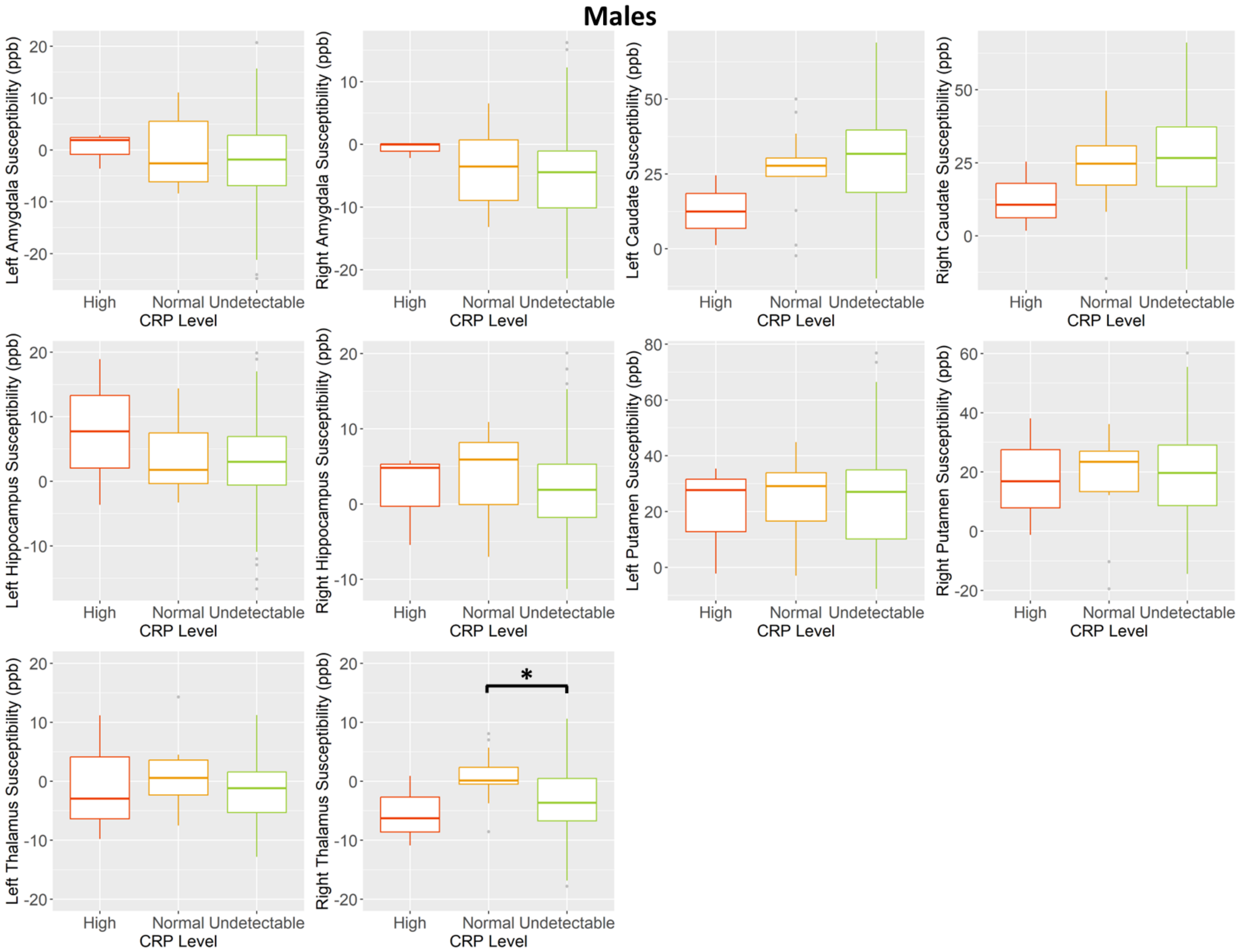
Males
Left Amygdala Susceptibility (ppb)
Right Amygdala Susceptibility (ppb)
Left Caudate Susceptibility (ppb)
Right Caudate Susceptibility (ppb)
Left Hippocampus Susceptibility (ppb)
Right Hippocampus Susceptibility (ppb)
Left Putamen Susceptibility (ppb)
Right Putamen Susceptibility (ppb)
Left Thalamus Susceptibility (ppb)
Right Thalamus Susceptibility (ppb)
High
Normal
Undetectable
CRP Level
*

Supplementary Figure 10. Box and whisker plots showing relationships in males between c-reactive protein (CRP) and iron in grey matter regions as measured by quantitative susceptibility mapping (n=155). High = >10mg/L, normal = 4-10mg/L, <4mg/L. * Indicates statistical significance before multiple comparison correction where $p<0.05$.

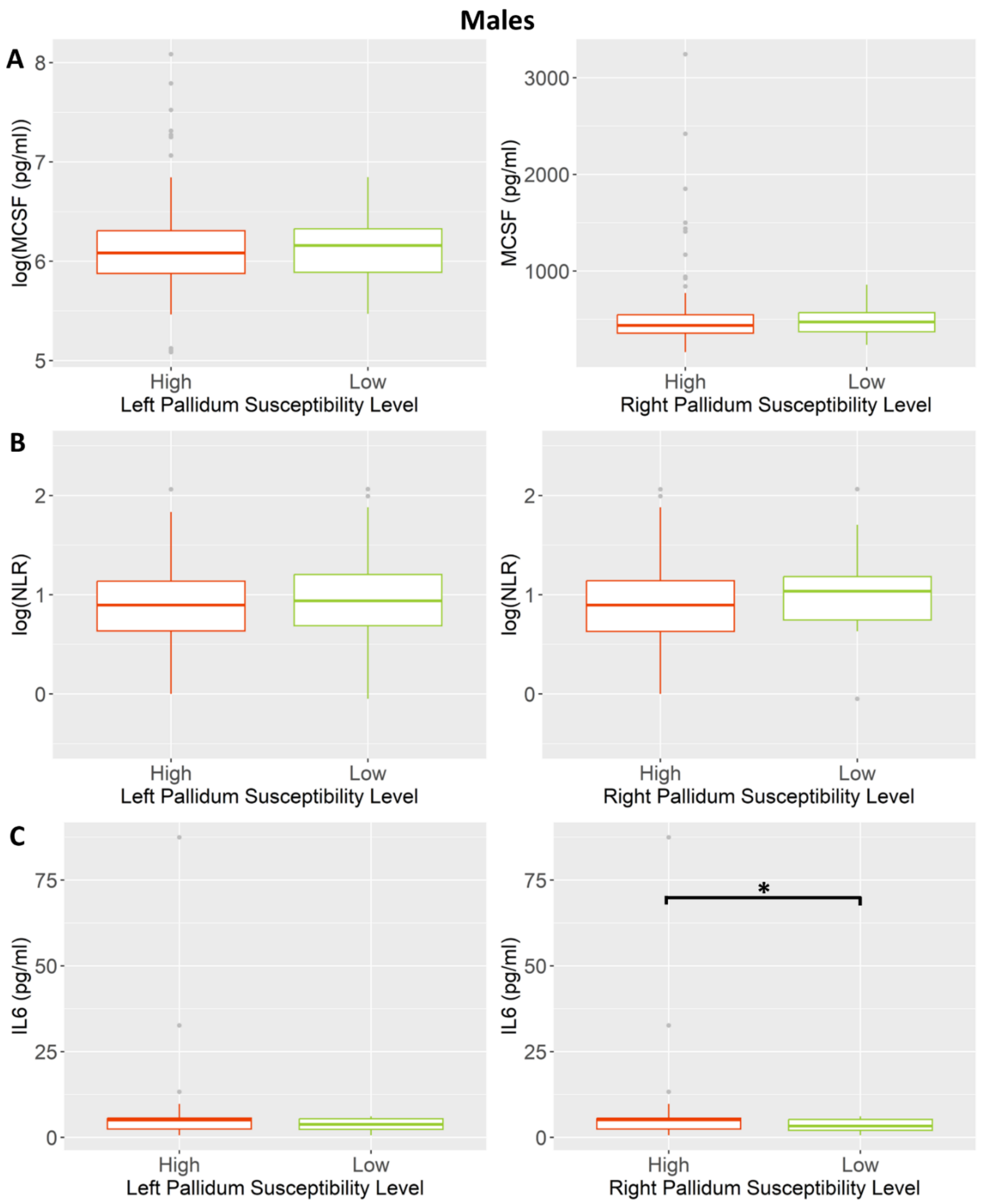


Supplementary Figure 11. A) Relationships in males between plasma macrophage colony stimulating factor 1 (MCSF) and brain iron in the pallidum as measured by quantitative susceptibility mapping (n=154). B) Relationships in males between neutrophil lymphocyte ratio and brain iron in the pallidum as measured by quantitative susceptibility mapping (n=155). C) Relationships in males between plasma interleukin – 6 (IL6) and brain iron in the pallidum as measured by quantitative susceptibility mapping (n=153). High = left pallidum susceptibility > 0.3ppm OR right pallidum susceptibility >0.125ppm, Low = left pallidum susceptibility < 0.3ppm OR right pallidum susceptibility < 0.125ppm. * Indicates statistical significance before multiple comparison correction where $p<0.05$.

**Males**

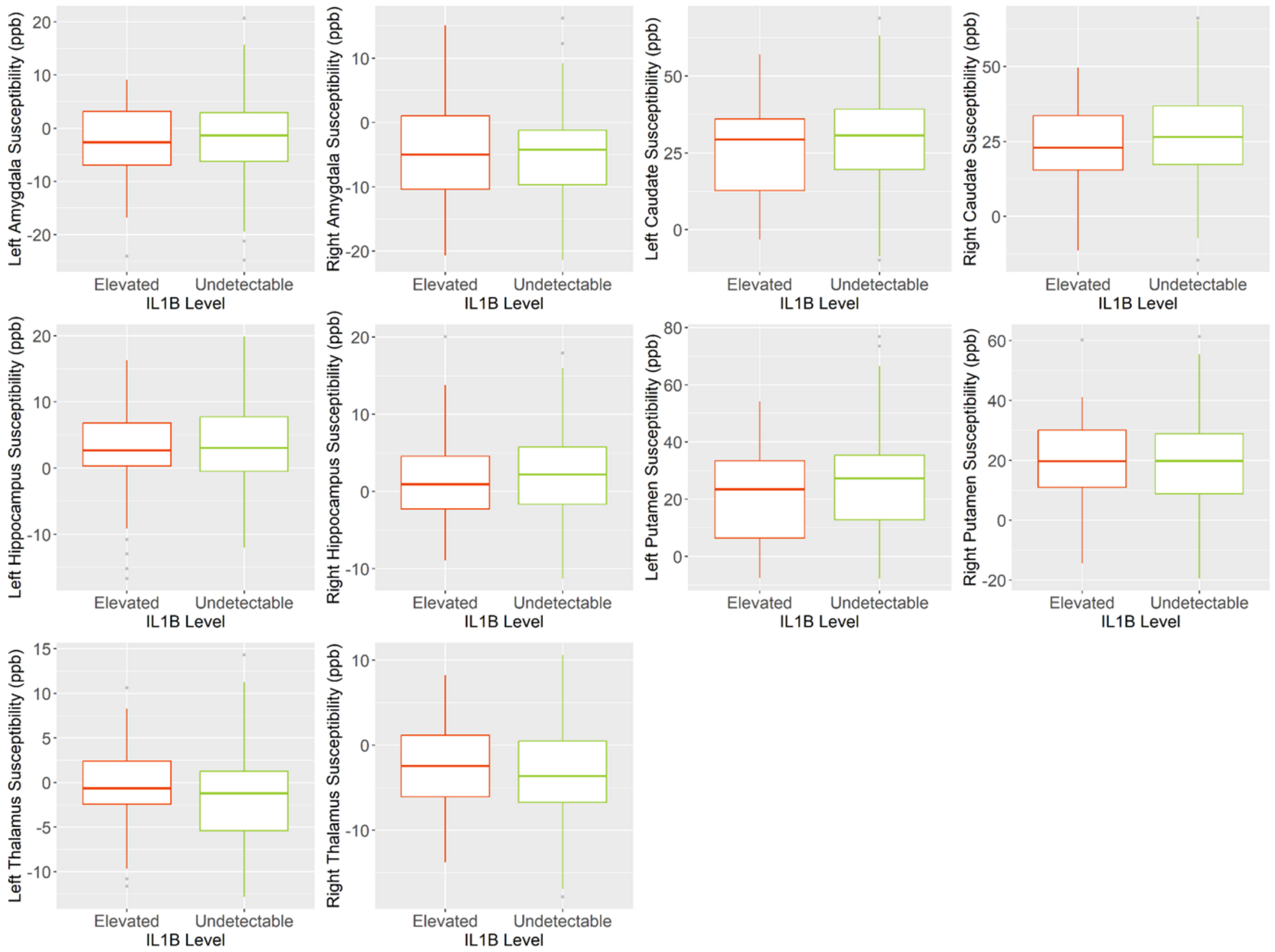

Left Amygdala Susceptibility (ppb)
Right Amygdala Susceptibility (ppb)
Left Caudate Susceptibility (ppb)
Right Caudate Susceptibility (ppb)
Left Hippocampus Susceptibility (ppb)
Right Hippocampus Susceptibility (ppb)
Left Putamen Susceptibility (ppb)
Right Putamen Susceptibility (ppb)
Left Thalamus Susceptibility (ppb)
Right Thalamus Susceptibility (ppb)
Elevated
Undetectable
IL1B Level

Supplementary Figure 12. Box and whisker plots showing the relationships in males between plasma interleukin 1 beta (IL1B) and iron in grey matter regions as measured by quantitative susceptibility mapping (n=151). Above = above detection limit (>0.92pg/ml), Below = below detection limit (<0.92pg/ml). * Indicates statistical significance before multiple comparison correction.
